# Inference of Self-Limiting Neutrophil Swarming Dynamics Using Bayesian Physics-Informed Neural Networks

**DOI:** 10.64898/2026.08.21.746187

**Authors:** Xincheng Wang, Pan Du, Karan Taneja, Jacob Doon-Ralls, Eduardo Reátegui, Maria A. Holland

## Abstract

Neutrophil swarming is a critical immune response in mammals and fish, in which neutrophils are recruited to inflammatory sites where they coordinate into a swarm that neutralizes pathogens. While excessive swarming can drive prolonged inflammation, a quantitative understanding of swarming dynamics remains limited. We developed a one-dimensional radial reaction–diffusion model of neutrophil swarming with two kinetic parameters, in order to capture the self-limiting swarming dynamics in both murine and human neutrophils in response to different inflammatory stimulus sizes. To ensure that the inverse problem is well-posed, we first performed sensitivity and identifiability analyses. We then developed a physics-informed neural network (PINN) to infer the key parameters governing swarm expansion and self-limitation. To account for uncertainty in noisy experimental measurements, we further extended this framework to a Bayesian PINN (B-PINN), which provides credible intervals for the inferred parameters. Both models were validated against synthetic data generated by numerical simulation and subsequently applied to *in vitro* experimental data from human and murine neutrophils in response to three bioparticle cluster sizes. The PINN-inferred dynamics show that larger bioparticle clusters are associated with greater cumulative recruitment and larger swarms in both species. The models further reveal species-specific differences in both the amplitude of initial recruitment and the timescale on which it self-limits. Additionally, the B-PINN posterior distributions quantify uncertainty in these species- and cluster size-dependent trends and identify where additional measurements would be most informative. To our knowledge, this is the first application of physics-informed machine learning to model neutrophil swarming dynamics. This framework provides a starting point for systematically comparing recruitment dynamics between human and murine neutrophils and offers guidance for future experimental design.

**Author Summary:** Neutrophils are small but among the most numerous immune cells, and among the first to respond to infection or tissue damage. They coordinate into dense clusters called swarms to isolate and destroy pathogens. Remarkably, swarming is self-limiting: recruitment eventually slows rather than continuing without control, helping protect healthy tissue from excessive inflammation. How these dynamics vary with stimulus size and between species remains difficult to quantify. Researchers can now trigger neutrophil swarming on a chip while precisely controlling the target size, but measurements of swarm growth are sparse and noisy. We combined a mathematical model of swarming with machine learning to infer these dynamics from such data. Our approach estimates the initial strength and duration of recruitment, with uncertainty bounds. Applying our approach to neutrophils from humans and mice, we found that larger targets drive stronger recruitment in both species, yet the two differ in the strength and timing of their response. Our analysis also showed that measuring the swarm radius early in swarm formation would be especially valuable for distinguishing these dynamics. This work provides a quantitative framework for designing more informative experiments, comparing swarming across species, and supporting translation of findings from mouse studies to human immune responses.

## 2 Introduction

Neutrophils are the most abundant immune cells in the human body. Around 10^11^ neutrophils are released from the bone marrow per day [1, 2]. With this massive number, they form the first line of the host defenses, where they respond rapidly to destroy pathogenic microorganisms through phagocytosis, degranulation, and the formation of extracellular traps [3]. In addition to their major roles in host defense, they also interact with other types of immune cells directly or by releasing cytokines, chemokines, and leukotrienes to modulate immune homeostasis.

Despite the variation across species and conditions, neutrophils display a remarkably conserved capacity for highly coordinated chemotaxis. When facing a threat, they roll and crawl along the endothelium and subsequently infiltrate into the interstitial tissue. After that, neutrophils follow a multi-step cascade and exhibit swarming behavior (Figure 1). The early short-range aggregation of pioneer neutrophils is triggered by pathogen/damage-associated molecular patterns (PAMPs/DAMPs) [4].

**Figure 1.**
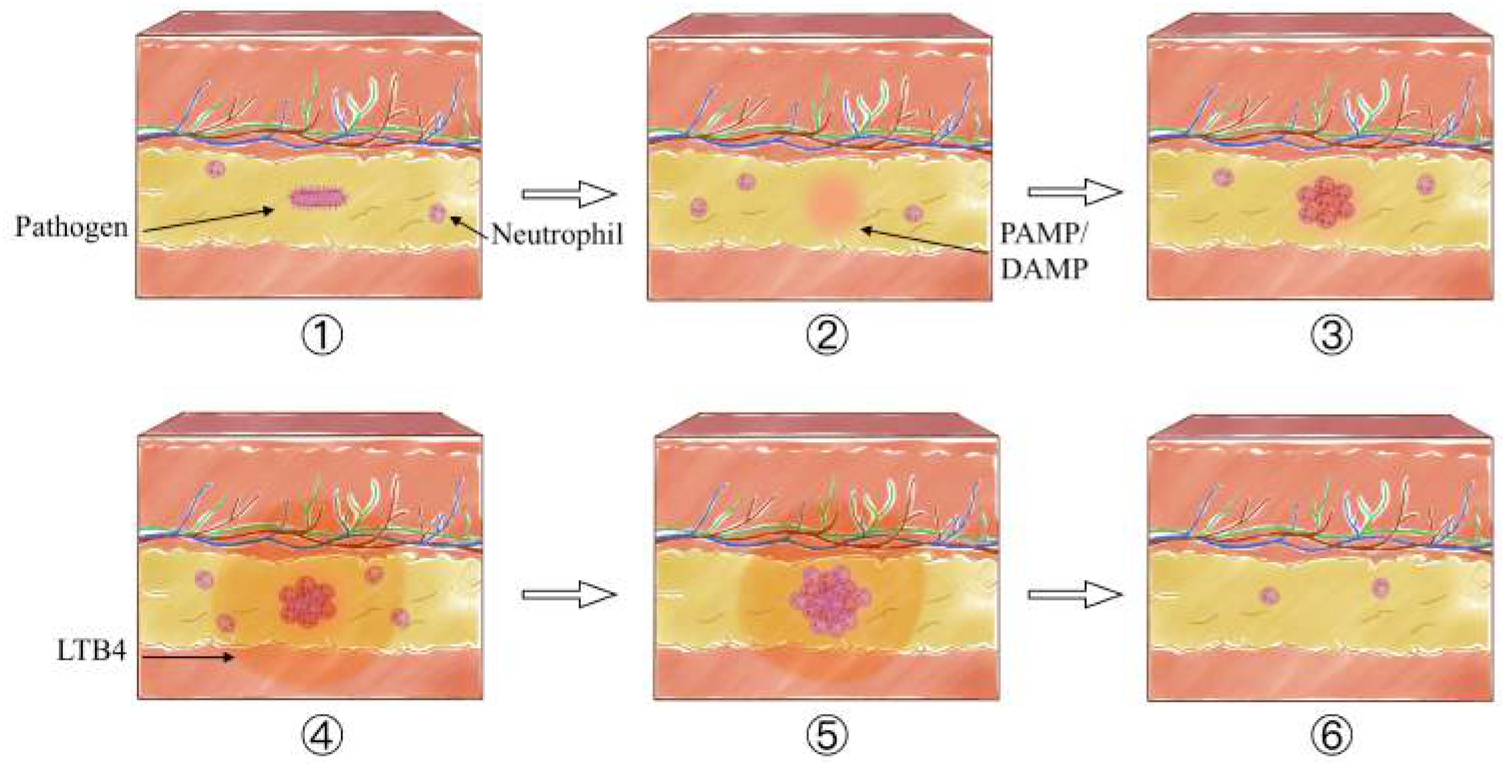
Neutrophil swarming follows a stepwise cascade. 1) A pathogen invasion or sterile tissue injury occurs in the tissue. 2) The pathogen or injury causes local cell death and the release of pathogen/damage-associated molecular patterns (PAMPs/DAMPs). 3) Pioneer neutrophils detect these signals and initiate the swarming, producing swarming-associated molecules, particularly LTB4. 4) LTB4 strongly amplifies swarming by recruiting additional neutrophils from the vasculature, and these newly recruited cells further raise the local LTB4 concentration, forming a positive feedback loop. 5) As the local concentration of LTB4 within the swarm reaches a threshold, G protein–coupled receptors (GPCRs) desensitize, which arrests neutrophil migration and prevents uncontrolled aggregation. 6) The pathogen is cleared and the swarm resolves.

When neutrophils form a cluster, they are activated and start producing swarming-associated molecules, particularly the lipid leukotriene B4 (LTB4). LTB4 amplifies the range of neutrophil recruitment [5, 6], as its chemical gradient reaches the vasculature [7]. This gradient recruits neutrophils continuously to the site [7] by encouraging long-range neutrophil vascular migration [8–10]. Recently, *in vivo* studies have also pointed out that calcium flux, connexin-43 (Cx43) hemichannels, and adenosine triphosphate (ATP) release are required for LTB4 production [11].

As more neutrophils migrate, a positive feedback loop is established, and the swarming is self-sustained [5]. However, the swarm does not expand or persist indefinitely. Although neutrophils continue to secrete chemoattractants throughout swarming, once the local concentration of the chemoattractant LTB4 exceeds a threshold, G protein–coupled receptors (GPCRs) on the neutrophils desensitize [12]. Desensitization halts further recruitment and transforms neutrophils from migrating hunters to stationary killers. This self-limiting regulatory mechanism prevents uncontrolled swarming, which could otherwise reduce phagocytic efficiency and lead to excessive inflammatory damage.

Beyond these molecular mechanisms, neutrophil swarming exhibits several robust quantitative features that hold across species and experimental systems. For example, swarming appears to require a critical threshold of local tissue damage to initiate: when the injury is minor, tissue-resident macrophages can ‘cloak’ the damage and prevent neutrophils from swarming [13], or if targets fall below a critical size threshold, individual neutrophils simply phagocytose the particles without triggering a collective swarm [14]. In other words, for small-scale tissue damage, it is easier and safer to handle it via other immune pathways. Once initiated, the size and geometry of the stimulus strongly affect the swarming response, and the final swarm size can differ roughly threefold between small and large bioparticle clusters.

While neutrophil swarms are an effective means of immobilizing and isolating pathogens, prolonged or excessive swarming can itself become harmful, causing collateral tissue damage, disrupting tissue homeostasis, and delaying tissue remodeling [3, 15]. Desensitization is a key mechanism of this protective swarm self-limiting. By disrupting the GPCR kinase signaling and thus desensitization, neutrophils move faster and spread out over larger areas, but they actually perform worse at bacterial elimination [12]. This demonstrates that desensitization leads to controlled arrest and makes swarming more effective. While the GPCR mechanism is important, it has also been shown that NADPH-oxidase-based negative feedback loop [16] contributes to the multi-factor process of swarm self-limitation.

Neutrophils exhibit several species-specific differences. In humans, neutrophils are the predominant leukocyte population, accounting for approximately 50–70% of circulating white blood cells [17], whereas in mice they represent only about 10–25% [18]. Human neutrophils also possess a relatively long circulatory half-life (approximately 5.4 days), whereas murine neutrophils survive for a much shorter period (approximately 12.5 hours) [18].

Despite these differences, mice and zebrafish are common animal models used to investigate swarming behavior. Mouse models provide unique advantages through transgenic reporter lines, enabling real-time *in vivo* tracking of neutrophil behavior [19]. Neutrophil swarming has been studied *in vivo* in mice at different tissue locations, including lungs, skin, liver, brain, and lymph nodes under both sterile and infectious conditions [5, 12, 20–24]. Transgenic lines of zebrafish larvae are also valued due to the ease of cell-specific green fluorescent visualization [11, 25–28]. However, direct translation of swarming findings from animal models to humans requires caution [29], and it is difficult to obtain physical measurements of swarming *in vivo* [30, 31]. *On the other hand, neutrophil microscale array experiments [12, 14, 32] provide an unprecedented capability to reproduce swarming in vitro*, enabling the precise control of stimulus size and location.

Partial differential equations (PDEs) have been utilized for years to analyze the dynamics of neutrophil signaling and chemotaxis [33–35]. Recently, neutrophil swarming in particular has been studied as a reaction–diffusion system using parabolic PDEs, whose traveling-wave solutions closely resemble the observed propagation of the swarm boundaries [36]. These models capture chemoattractant diffusion in different dimensions [36] and the wave-like dynamics of neutrophil activator and inhibitor molecules [16]. However, these models focus on molecular-level diffusion and pay less attention to cell movement, partly because of the lack of experimental cell counting and the difference in velocities between chemoattractants and cells [16].

Machine learning has become an increasingly important tool for discovering and calibrating dynamical systems in biomedical science [37–40]. Among these approaches, physics-informed neural networks (PINNs) provide a hybrid framework that combines governing physical laws with available experimental data, and have been used for parameter estimation and inverse modeling [41–43]. For example, Zhang et al. [44] combined PINNs with symbolic regression to discover a reaction–diffusion model for misfolded tau protein spreading in Alzheimer’s disease from clinical imaging data, demonstrating the potential of PINNs for learning interpretable biomedical dynamics. However, experimental measurements are often sparse and noisy, making it essential not only to infer model parameters but also to quantify uncertainty in those estimates. Bayesian inference is a principled framework for addressing this need [45], and B-PINNs in particular have been used to quantify parameter uncertainty across a range of nonlinear dynamical systems [46–48]. Despite these advances, machine-learning-based inverse modeling remains underexplored in neutrophil swarming, where experimental data are typically limited by sparse temporal sampling, replicate-to-replicate variability, and measurement noise. Previous experimental and mathematical studies have provided important insight into neutrophil chemotaxis and swarming dynamics [14, 16, 49–52], but PINN- and B-PINN-based frameworks have not yet been developed to infer interpretable recruitment parameters and quantify their uncertainty from swarm-front observations.

The experimental data analyzed in this work come from a recent *in vitro* swarming study. To gain insights into subcellular processes, Glaser et al. [19] combined the swarming-on-a-chip platform with confocal laser-scanning microscopy to investigate two distinct regulatory pathways during the later stages of swarming. In this setup, arrays of heat-killed *Staphylococcus aureus* (HKSA) bioparticles were patterned into circular clusters with different sizes on glass coverslips, mimicking local infection and triggering neutrophils to release LTB4 and initiate swarming. This platform enabled high-resolution, explicit quantification of swarming cell numbers in both human and murine neutrophils. Interestingly, not only did the bioparticle cluster size significantly influence the final swarm area, but the two species also exhibited different swarming speeds under the same cluster-size conditions. However, the kinetic parameters governing these bioparticle cluster size- and species-dependent differences have not yet been quantified.

Despite growing experimental and computational studies of neutrophil swarming, several gaps remain. First, can a continuum model reproduce the wave-like propagation of neutrophil recruitment and the self-limiting behavior of the swarm front propagation? Second, can such a model recover interpretable kinetic parameters that quantify how swarming depends on the size of the inflammatory stimulus? Third, can the framework identify species-level differences between human and murine neutrophils while accounting for the uncertainty introduced by sparse, noisy measurements? To address these questions, we model neutrophil swarming as a one-dimensional radial reaction–diffusion system with a time-dependent recruitment rate that encodes GPCR desensitization (section 3). Before inverse modeling, we perform sensitivity and identifiability analyses to confirm that the inverse problem is well-posed and to identify which parameters are most reliably recovered from swarm-front trajectory data (subsection 4.1). We then develop a PINN to infer the governing recruitment parameters and validate it on perfect synthetic data (subsection 4.2). To quantify uncertainty in the inferred parameters under noisy experimental measurements, we extend the framework to a B-PINN that provides posterior distributions and credible intervals (subsection 4.3). Finally, we apply both models to *in vitro* human and murine neutrophil swarming data across three bioparticle cluster sizes, and report the resulting species- and size-dependent recruitment trends together with their uncertainty (subsection 4.4, subsection 4.5).

## 3 Methodology

### 3.1 Governing equation

In this section, we assume a continuum description of neutrophil swarming. We first adopt a typical reaction-diffusion equation for the density of neutrophils at a point,

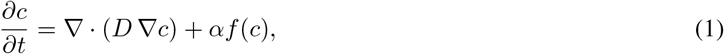

where *c* ∈ [0, 1] is the dimensionless normalized neutrophil density at a point, defined as *c* = *C/C*_max_ with *C* the local areal cell density and *C*_max_ the maximum carrying capacity; *D* [µm^2^ min^−1^] characterizes the effective cell motility; *α* [min^−1^] is the recruitment rate constant; and *f* : ℝ → ℝ is the source term. Here, the first term on the right-hand side represents the tendency of cells to spread out (diffusion), while the second term represents the recruitment of additional neutrophils.

The primary goal of this study is to identify an appropriate form for the second term such that the equation captures the spatial and temporal distribution of neutrophils. For the source term, we adopt the well-studied Fisher–Kolmogorov–Petrovskii–Piskunov (Fisher–KPP) form, *f*(*c*) = *c*(1 − *c*) [53], derived from logistic growth. This form captures three key aspects of neutrophil swarming: (i) without an initial population of pioneer neutrophils (*c* = 0), swarming cannot be initiated [54]; (ii) at full local occupancy (*c* = 1), no further cells can be recruited; and (iii) it admits traveling-wave solutions that propagate at constant speed while preserving wavefront shape, analogous to the observed evolution of neutrophil swarm boundaries. However, the self-extinguishing behavior of neutrophil swarms is not captured by autonomous Fisher-KPP models [36, 55, 56].

To reproduce the experimentally observed shutting down of the swarm expansion at later times, we modify the rate constant *α* into a time-dependent function *α*(*t*)

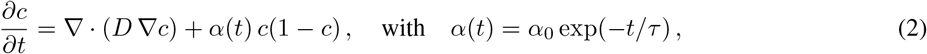

where *α*(*t*) is the recruitment rate function and is assumed to decrease monotonically. This phenomenologically represents the gradual attenuation of neutrophil recruitment associated with LTB4 receptor desensitization. It has two parameters: *α*_0_ [min^−1^] sets the initial recruitment amplitude, driven by the initial sensitivity, while the characteristic timescale *τ* [min], analogous to the relaxation time in viscoelastic Maxwell models, governs the timescale of receptor desensitization and therefore the slowing of recruitment. Experimental observations show nonlinear evolution of the swarm front [12, 14], motivating this exponential decay form. As lim_*t*_→∞ *α*(*t*) = 0, receptor signaling becomes completely suppressed, halting further neutrophil recruitment and effectively terminating the swarming process. The model variables and parameters used in this study are summarized in Table 1.

**Table 1.** Model variables, parameters, experimental conditions, and numerical settings used in this study. PINN and B-PINN settings specific to each species and bioparticle cluster size are reported separately in Table 2 and Table 3.

| Description | Symbol | Value & units |
| --- | --- | --- |
| *State variable and coordinates* |  |  |
| Normalized neutrophil density | $c$ | dimensionless, $\in [0, 1]$ |
| Radial coordinate | $r$ | 0 to 100 $\mu\text{m}$ |
| Time | $t$ | 0 to 120 min |
| Swarm-front radius | $R(t)$ | determined ($\mu\text{m}$) |
| Terminal swarm radius | $R(t_{\max})$ | determined ($\mu\text{m}$) |
| *Experimental conditions* |  |  |
| HKSA bioparticle cluster diameter | — | 30 $\mu\text{m}$, 60 $\mu\text{m}$ and 120 $\mu\text{m}$ |
| *PDE model parameters* |  |  |
| Effective motility (diffusion) coefficient | $D$ | 0.1 $\mu\text{m}^2 \text{min}^{-1}$ |
| Length scale of initial Gaussian profile | $R_0$ | 10 $\mu\text{m}$ |
| Initial recruitment amplitude | $\alpha_0$ | inferred ($\text{min}^{-1}$) |
| Recruitment decay timescale | $\tau$ | inferred (min) |
| Cumulative recruitment measure | $\beta = \alpha_0 \tau$ | derived (dimensionless) |
| *Numerical discretization* |  |  |
| Number of radial grid nodes | $N$ | 101 |
| Temporal collocation points (PDE residual) | $N_{\text{col}}$ | 500 |

Swarming experiments are typically conducted in two-dimensional domains. Assuming radial symmetry of the neutrophil cluster (Figure 3), we express the neutrophil density as *c* = *c*(*r, t*) where *r* is the radial coordinate, reducing the governing equation to a one-dimensional radial problem:

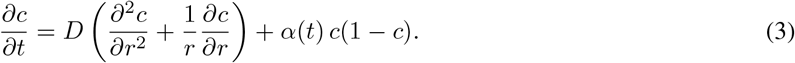

To reduce the complexity of the learning problem and avoid the computational cost of repeatedly evaluating second-order spatial derivatives via automatic differentiation [44, 57], we construct a discrete radial Laplacian operator ***L*** using a second-order finite-difference scheme. This converts the PDE into a system of ordinary differential equations (ODEs) on a radial grid of *N* = 101 points. The semi-discrete form of Equation 3 is:

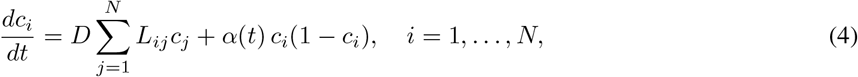

where *L*_*ij*_ are the entries of the discrete radial Laplacian operator.

Reaction–diffusion systems are highly sensitive to initial conditions, which significantly influence the subsequent traveling-wave dynamics. To ensure consistency in inferred results, we prescribe a Gaussian initial profile centered at the origin to approximate the experimentally observed initial accumulation of pioneer neutrophils:

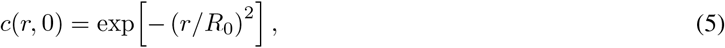

where *R*_0_ = 10 µm is the length scale of the initial Gaussian profile. This smooth initialization supports the rapidly growing phase of the swarm while preserving wavefront shape.

We define the computational domain as *r* ∈ [0, 100 µm]. A symmetry boundary condition is imposed at the origin to ensure regularity of the radial solution, while a zero-flux Neumann boundary condition is applied at the outer radius.

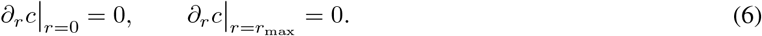

### 3.2 Synthetic data preparation

To validate our inference frameworks, we first generated synthetic data by solving the governing equation (Equation 3) with prescribed parameters. The diffusion coefficient was set to *D* = 0.1 µm^2^ min^−1^ to represent slow and contact-mediated motility of swarming neutrophils. It also allowed us to model the sharp, non-dispersive wavefront observed in experiments [19]. The recruitment parameters were prescribed as *α*_0_ = 1.0 min^−1^ and *τ* = 10 min. The swarm-front radius *R*(*t*) is defined as the radial coordinate *R* at which *c*(*R, t*) = 0.5.

The governing PDE was solved numerically using `SciPy`, and the resulting density profiles were sampled at *N*_obs_ = 7 discrete observation times, *t*_*i*_ ∈ {0, 10, 20, 30, 60, 90, 120} min. As our focus for the PINN validation is to confirm that the parameters can be retrieved even for sparse data, we constructed a noise-free synthetic dataset for PINN training (Figure 2, left); the corresponding training settings are listed in Table 2.

**Table 2.** PINN training settings used for synthetic and experimental datasets. For all PINN fits, the PDE residual was evaluated on *N*_col_ = 500 dense temporal collocation points, with 10^5^ Adam [61] iterations followed by 10^4^ L-BFGS [62] iterations.

| Dataset | Condition | $\lambda_{\text{WD}}$ | $s_{\text{max}}$ | $\delta$ |
| --- | --- | --- | --- | --- |
| Synthetic | – | 0 | 1 | – |
| Human | 30 $\mu\text{m}$ | $10^{-6}$ | 1 | 0.100 |
| Human | 60 $\mu\text{m}$ | $10^{-6}$ | 300 | 0.100 |
| Human | 120 $\mu\text{m}$ | $10^{-6}$ | 1 | 0.100 |
| Murine | 30 $\mu\text{m}$ | $10^{-7}$ | 100 | 0.005 |
| Murine | 60 $\mu\text{m}$ | $10^{-7}$ | 10 | 0.005 |
| Murine | 120 $\mu\text{m}$ | $10^{-7}$ | 10 | 0.005 |

**Figure 2.**
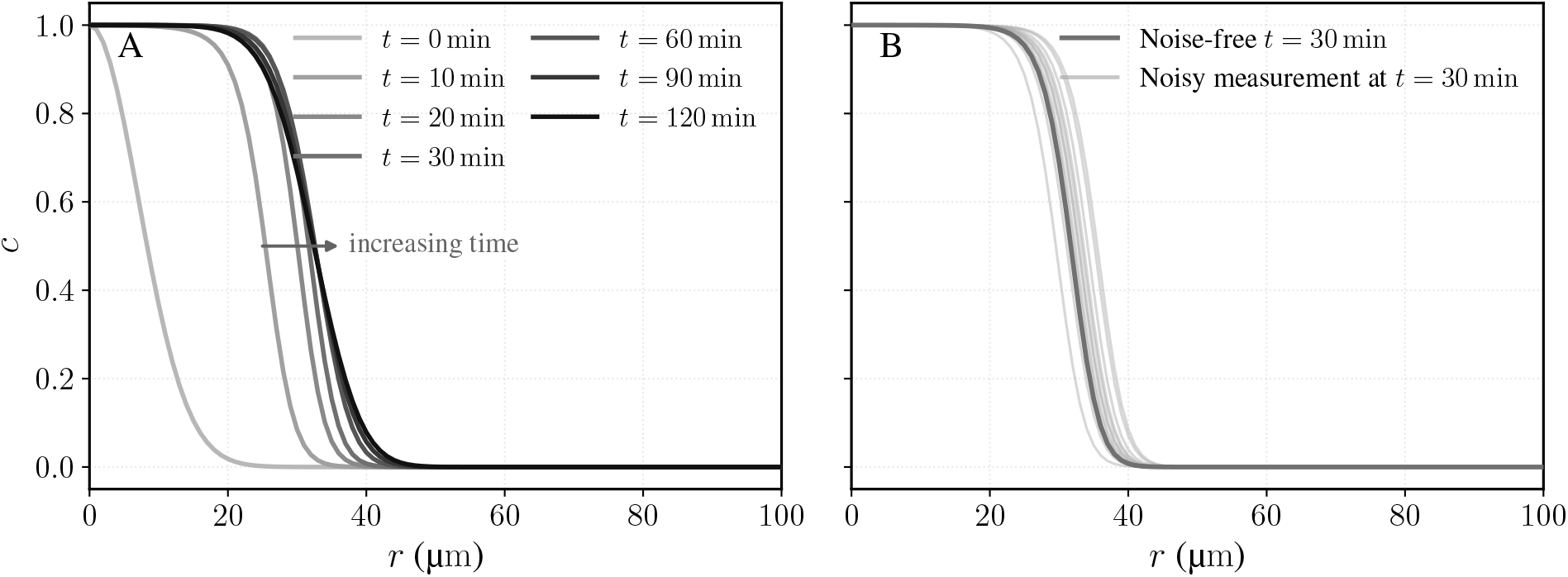
Synthetic normalized neutrophil density profiles generated from the numerical solution of the governing PDE. (A) Noise-free concentration profiles *c*(*r, t*_*i*_) at *t*_*i*_ ∈ {0, 10, 20, 30, 60, 90, 120} min, used for PINN training. (B) Representative noisy normalized neutrophil density profiles at *t* = 30 min; for visualization, the ground-truth density profile (black line) is plotted together with a min–max envelope computed from the noisy replicates at each radial position (shaded bands). These noisy realizations illustrate the measurement variability used for B-PINN training.

For the subsequent B-PINN analysis, we generated *N*_exp_ = 12 noisy synthetic replicates from the same ground-truth profiles by perturbing the apparent wavefront location (Figure 2, right). Specifically, at each observation time *t*_*i*_ *>* 0, the profile *c*(*r, t*_*i*_) was shifted by a random displacement 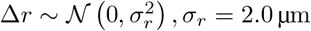 µm, and then resampled on the original radial grid using cubic interpolation. This perturbation simulates uncertainty in the measured swarm-front location while preserving the shape of the radial density profile. The initial condition at *t* = 0 was left unperturbed and was included identically across all synthetic replicates.

### 3.3 Experimental data preparation

To evaluate the biological relevance of our modeling approach, we used the human and murine neutrophil swarming datasets reported by Glaser et al. [19] (Figure 3). Swarming assays were conducted on an adapted swarming-on-a-chip platform, where neutrophils were exposed to immobilized HKSA bioparticle clusters with diameters of 30 µm, 60 µm, and 120 µm. Both human and murine neutrophils exhibited self-extinguishing swarming behavior, although their migration speeds and final swarm sizes differed across species and bioparticle cluster sizes.

Swarm size was quantified experimentally by manually counting the number of neutrophils within each swarm over time. For each HKSA bioparticle cluster diameter, three independent trials were performed. In the human neutrophil dataset, four samples were analyzed per trial, yielding *N*_exp_ = 12 independent replicates per condition, each recorded at *N*_obs_ = 7 observation times, *t*_*i*_ ∈ {0, 10, 20, 30, 60, 90, 120} min. In the murine dataset, four to six samples were analyzed per trial, giving *N*_exp_ = 15–18 replicates per condition (Table 3), each recorded at *N*_obs_ = 9 observation times, *t*_*i*_ ∈ *{*0, 10, 20, 30, 40, 50, 60, 90, 120*}* min.

**Table 3.** Likelihood noise scales and number of independent replicates used for each B-PINN inference. The synthetic case uses constant likelihood noise scales, whereas the experimental likelihood noise scales were selected separately for each species and bioparticle cluster size to account for differences in replicate measurement variance. For all conditions, the log-normal prior variances were fixed at 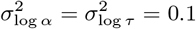 and the network-weight prior standard deviation at *σ*_*θ*_ = 0.02.

| Dataset | Condition | $N_{\text{exp}}$ | $\sigma_c$ | $\sigma_R$ |
| --- | --- | --- | --- | --- |
| Synthetic | – | 12 | 0.020 | 0.200 |
| Human | 30 $\mu\text{m}$ | 12 | 0.010 | 0.150 |
| Human | 60 $\mu\text{m}$ | 12 | 0.010 | 0.200 |
| Human | 120 $\mu\text{m}$ | 12 | 0.010 | 0.300 |
| Murine | 30 $\mu\text{m}$ | 15 | 0.015 | 0.100 |
| Murine | 60 $\mu\text{m}$ | 18 | 0.010 | 0.500 |
| Murine | 120 $\mu\text{m}$ | 17 | 0.020 | 1.000 |

To estimate the size of the swarm from the measured cell counts, we used a geometric approximation. Each neutrophil was modeled as a circular cell with radius *r*_cell_ = 5 µm [19]. We approximated the swarm as a tightly packed disk in which the total projected area equals the sum of individual cell footprints, 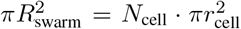, where *N*_cell_ denotes the counted number of neutrophils in the swarm. Under this assumption, the maximum carrying capacity is 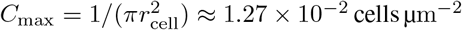 cells µm^−2^, and the swarm radius is

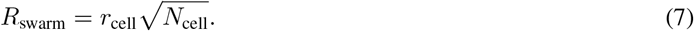

From this calculated swarm radius, we reconstructed an approximate radial cell density profile for PINN and B-PINN training. Specifically, we used numerically generated reference profiles described in subsection 3.2 and shifted each profile in the radial direction so that the location where *c* = 0.5 matched the experimentally inferred swarm radius *R*_swarm_ at the corresponding time point. For PINN training on the experimental data (Figure 4), we averaged all samples across trials at each measurement time, denoted by 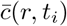. The corresponding radial cell profiles were then reconstructed using the method described in subsection 3.2. While this approach assumes certain values for the parameters, the swarm profile tends to retain a similar wavefront shape for a reasonable range of parameters, so the full radial field can be approximated given only the swarm radius.

**Figure 3.**
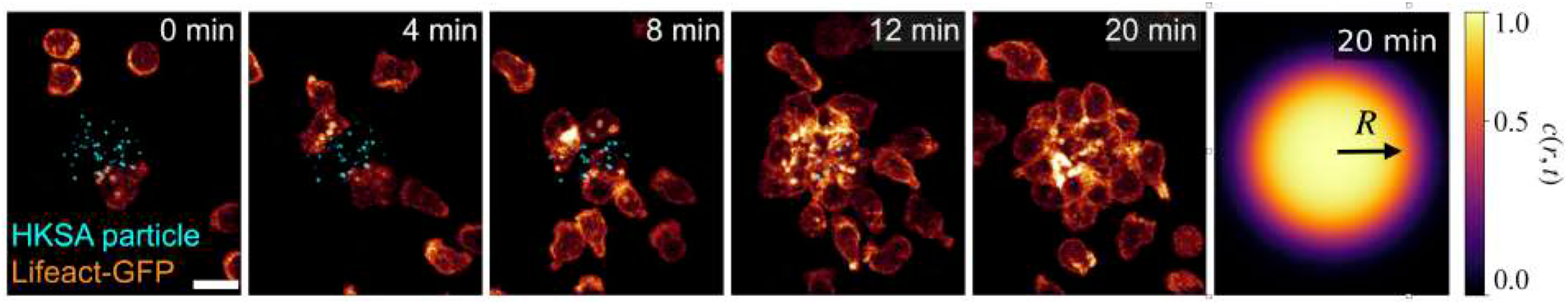
Experimental time series of murine neutrophil swarming and an illustrative model snapshot. The first five panels show murine neutrophils expressing Lifeact-GFP, a fluorescent reporter of filamentous actin, as they swarm toward an immobilized HKSA bioparticle cluster (cyan) at 0, 4, 8, 12, and 20 minutes. The glow heatmap represents Lifeact-GFP fluorescence intensity. Scale bar: 10 µm. The rightmost panel shows a top-view snapshot of the normalized neutrophil density *c*(*r, t*) generated by the radially symmetric Fisher–KPP model at *t* = 20 min. The arrow denotes the instantaneous swarm-front radius *R*(*t*). Experimental images were adapted from Figure 1D of Glaser et al. [19] under the Creative Commons Attribution 4.0 International license (https://creativecommons.org/licenses/by/4.0/).

**Figure 4.**
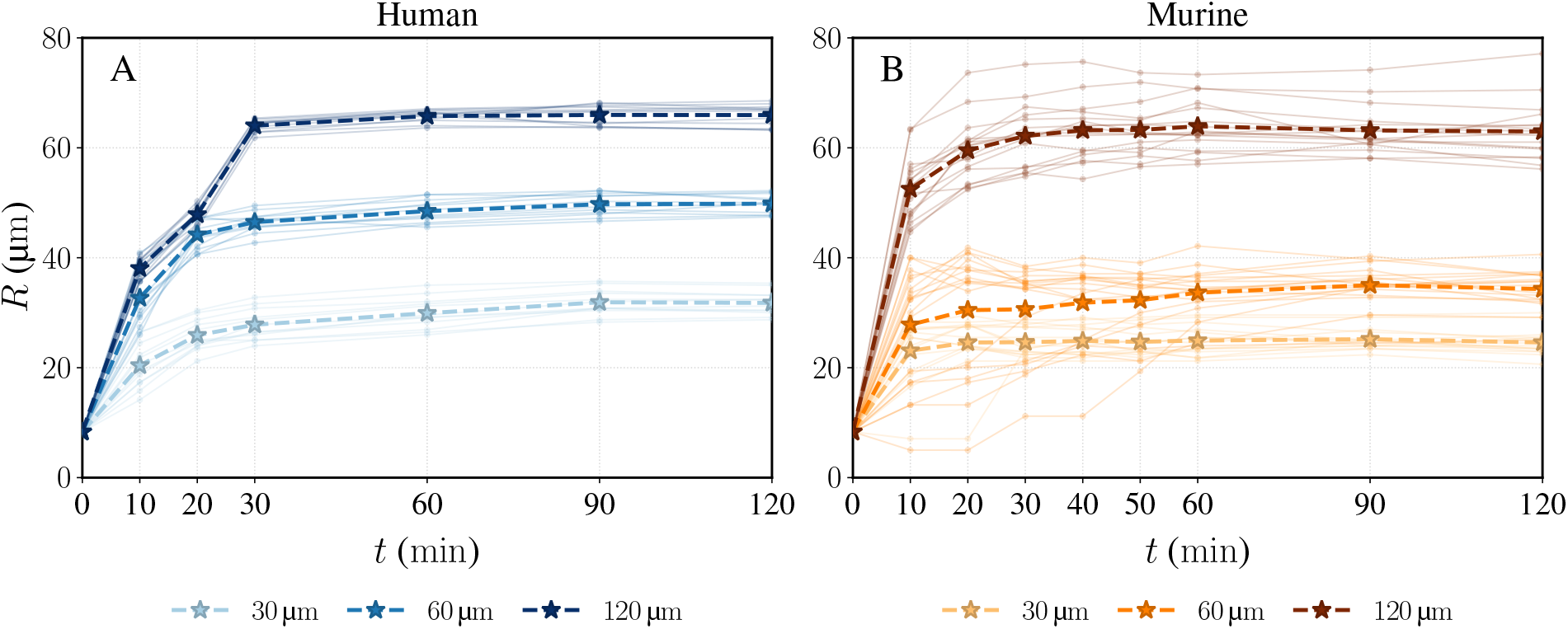
Experimental swarm-front trajectories for human (A) and murine (B) neutrophils across three bioparticle cluster sizes.

**Figure 5.**
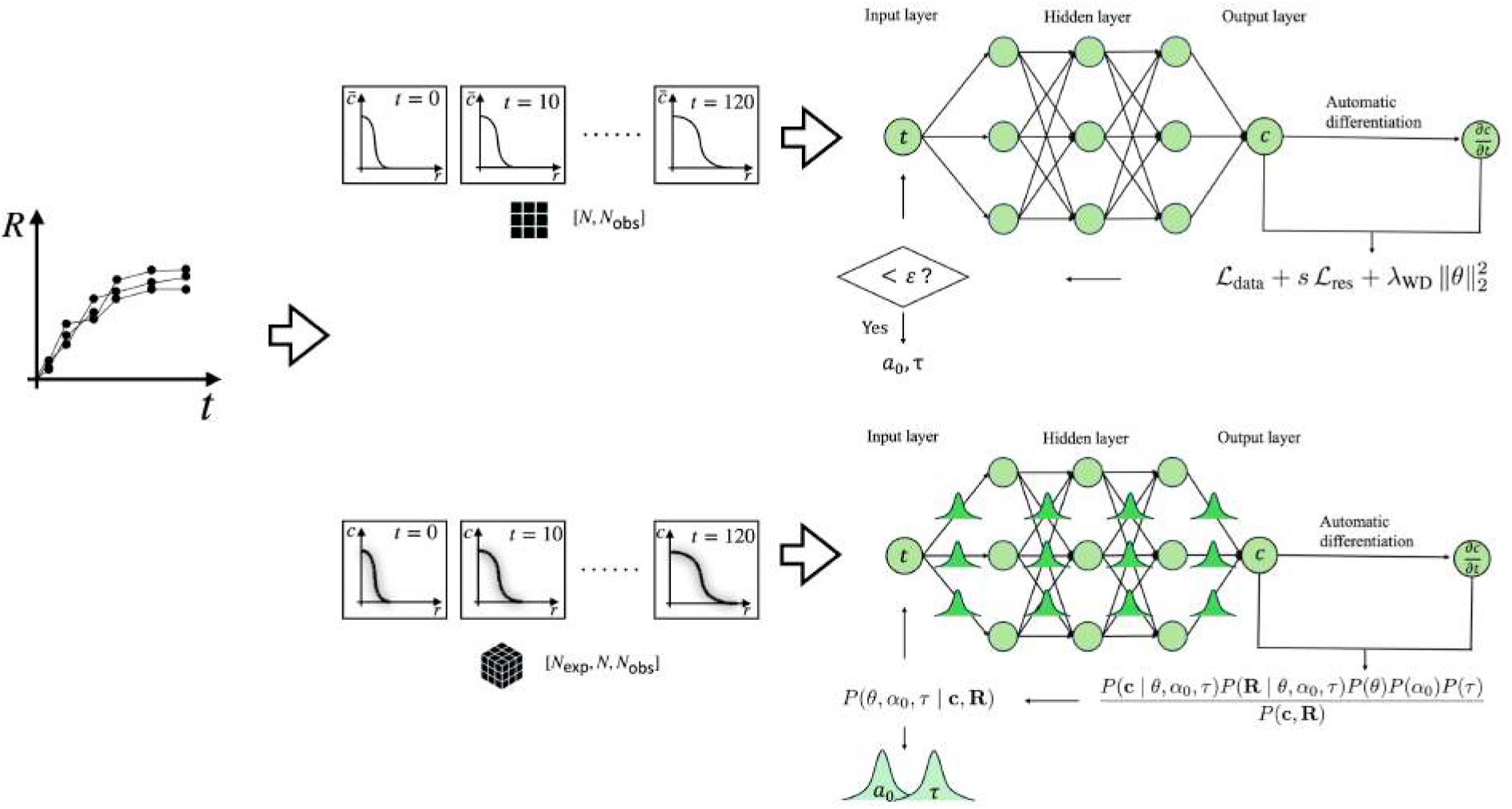
Schematic of the PINN (top) and B-PINN (bottom) frameworks. The networks take time *t* as input and output the full radial field ***ĉ***_*θ*_(*t*). The prediction and its temporal derivative *d****ĉ***_*θ*_*/dt* are used to compute the total loss ℒ_total_, defined as shown. The recruitment parameters *α*_0_ and *τ*, or in the case of the B-PINN, their distributions, are inferred jointly with the network parameters *θ*.

### 3.4 Physics-Informed Neural Networks

Having established the governing model and prepared the synthetic and experimental data, we now describe the inference frameworks, beginning with the deterministic PINN [58] used to jointly learn the cell density field *c*(*r, t*) and infer the unknown recruitment parameters *α*_0_ and *τ* from sparse experimental observations. The PINN is a multilayer perceptron ***ĉ***_*θ*_(*t*) with parameters *θ* = {***W*** _*k*_, ***b***_*k*_*}* that maps a scalar time input *t* to the full radial density at all *N* grid nodes simultaneously, ***ĉ***_*θ*_(*t*) ∈ ℝ^*N*^ [44] (Figure 5). The network has six hidden layers of 50 neurons each with hyperbolic tangent (tanh) activation and is implemented in JAX. The recruitment parameters *α*_0_ and *τ* are treated as trainable variables alongside the network parameters θ, so that all dim(θ) + 2 parameters are optimized jointly.

The training process minimizes a total loss that combines a data-misfit term ℒ_data_, a PDE-residual term ℒ_res_, and an *L*_2_ weight-decay regularizer:

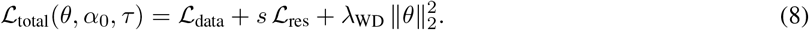

The data-misfit term ℒ_data_ enforces agreement between the network predictions and experimental observations at *N*_obs_ sparse time points. To reduce the sensitivity of the PINN to measurement outliers, we adopt the Huber loss [59] with threshold δ:

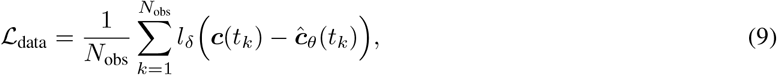

where ***c***(*t*_*k*_) = [*c*_1_(*t*_*k*_), …, *c*_*N*_ (*t*_*k*_)]^⊤^ ∈ ℝ^*N*^ is the vector of normalized cell density at the *N* radial nodes at time *t*_*k*_, ***ĉ***_*θ*_ is the corresponding network prediction, and *l*_*δ*_ is the Huber loss applied elementwise and averaged over components:

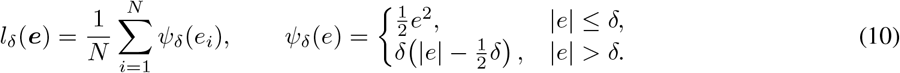

The physics term penalizes the discrepancy between the network time derivative and the right-hand side of the semi-discrete governing equation. At any time *t*, we define the vector-valued PDE residual as

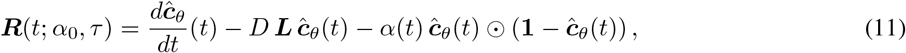

where ***R***(*t*; *α*_0_, *τ*) ∈ ℝ^*N*^, ***L*** ∈ ℝ^*N ×N*^ is the discrete radial Laplacian introduced in Equation 4, ⊙ denotes the elementwise product, and **1** ∈ ℝ^*N*^ is a vector of ones.

The PDE residual loss ℒ_res_ averages the squared residual over both the *N*_col_ temporal collocation points and the *N* radial components:

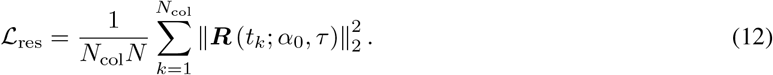

The data-misfit and PDE-residual losses have substantially different magnitudes, which can cause one term to dominate the gradient signal [60]. To address this, we rescale the PDE-residual loss using a lightweight adaptive factor:

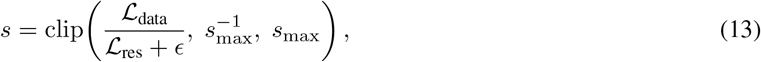

with *ϵ* = 10^−10^ for numerical stability. The scale factor *s* is treated as a stop-gradient quantity, keeping the data and residual contributions to the total loss on comparable scales throughout training. The weight-decay coefficient *λ*_WD_, clipping threshold *s*_max_, and Huber threshold δ were selected separately for each experimental condition (Table 2).

The data loss is evaluated at the experimental observation times, while the PDE residual is enforced at dense temporal collocation points over the full simulation interval. Training proceeds in two phases, using Adam optimization followed by L-BFGS refinement to ensure convergence.

### 3.5 Bayesian Physics-Informed Neural Networks

Unlike the PINN framework introduced above, which infers deterministic parameter values but does not quantify uncertainty, Bayesian inference incorporates both measurement data and physics constraints through probabilistic likelihood models [46]. For noise-free synthetic data, PINNs accurately recover the ground-truth parameters. In practical settings, however, experimental observations are contaminated by measurement noise, and the inferred parameters are subject to uncertainty arising from noisy and limited observations.

Bayesian inference quantifies this uncertainty by combining likelihoods associated with the observed cell density data ***c*** and the physics residual ***R*** with prior distributions over the model parameters, yielding a posterior distribution over *θ, α*_0_, and *τ* . We integrate the Bayesian framework with the PINN model to infer these posteriors and propagate the resulting uncertainty through the reaction–diffusion equation, yielding credible intervals for the predicted swarming dynamics. Using Bayes’ theorem and assuming that the data and physics-residual likelihoods are conditionally independent and that the priors on *θ, α*_0_, and *τ* are mutually independent, the joint posterior factorizes as

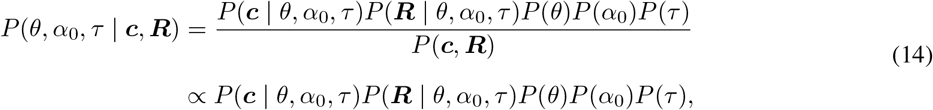

where *P*(***c*** |*θ, α*_0_, *τ*) and *P*(***R*** |*θ, α*_0_, *τ*) are the data and physics-residual likelihoods, respectively, and *P*(*θ*), *P*(*α*_0_), and *P*(*τ*) are the prior distributions of the model parameters.

The data likelihood quantifies the agreement between the prediction ***ĉ***_*θ*_(*t*) and the observed data ***c***(*t*) at the discrete observation times *t*_*i*_. Assuming independent Gaussian measurement noise with variance 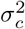, it factorizes over all scalar observations,

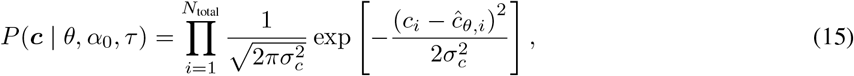

where *N*_total_ = *N*_exp_ × *N*_obs_ × *N* is the total number of scalar observations, *N*_exp_ is the number of independent replicates used for each inference (Table 3), *c*_*i*_ and ĉ_*θ,i*_ are the *i*-th entries of the flattened observation and prediction vectors, indexed jointly over the total number of scalar observations, and *σ*_*c*_ is the likelihood noise scale (Table 3).

The physics-informed likelihood evaluates how accurately the prediction ***ĉ***_*θ*_(*t*) satisfies the governing PDE. Assuming independent Gaussian residual noise with variance 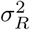, the physics-informed likelihood is:

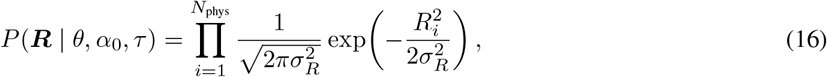

where *N*_phys_ = *N*_obs_ *× N* is the number of scalar residual entries, *R*_*i*_ denotes the *i*th entry of the flattened residual vector ***R***, indexed over observation time and radial position, and *σ*_*R*_ is the likelihood noise scale (Table 3).

The terms *P*(*θ*), *P*(*α*_0_), and *P*(*τ*) denote the prior distributions of the network parameters *θ* and the recruitment parameters *α*_0_ and *τ*, respectively. The PINN solution is used both as the initial state for Hamiltonian Monte Carlo (HMC) sampling and as the center of the prior distribution. Specifically, we assign a Gaussian prior to the network parameters centered on the PINN weights, and we impose log-normal priors on the recruitment parameters to ensure positivity. Equivalently, Gaussian priors are placed on the log-transformed parameters, which are mapped back to physical space during sampling.

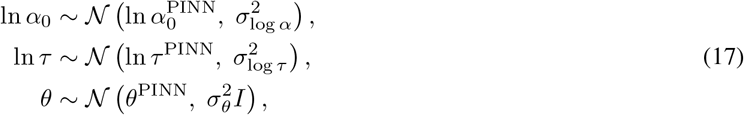

where 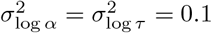 and *σ*_*θ*_ = 0.02.

To infer the posterior distributions, we used the No-U-Turn Sampler (NUTS) [63], an adaptive extension of HMC [64, 65], as implemented in NumPyro [66]. We ran four chains, each with 1000 warm-up iterations followed by 2000 retained samples, at a target acceptance probability of 0.8 and a maximum tree depth of 12. During warm-up, the step size and mass matrix were adapted. The mass matrix used a dense block for the two log-transformed recruitment parameters (ln *α*_0_, ln *τ*) and a diagonal structure for the neural-network parameters *θ*. Chains were initialized at dispersed points around the deterministic PINN estimate. Sampling quality was assessed using the effective sample size (ESS), the potential scale reduction factor 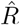, and the number of divergent transitions.

To propagate parameter uncertainty into the predicted swarm dynamics, we solved the forward PDE for an ensemble of 500 parameter pairs drawn from the HMC posterior. The posterior predictive mean was computed as the ensemble mean and the 95% predictive band was bounded by the 2.5th and 97.5th percentiles of the resulting trajectories.

## 4 Results

### 4.1 Analysis of the forward and inverse modeling framework

Before pursuing inverse inference of the swarming parameters, we examine whether the inverse problem is *well-posed* [67]; that is, whether (i) a solution exists, (ii) the solution is unique, and (iii) the solution depends continuously on the data. We assess these conditions through a parametric sensitivity study, an asymptotic analysis of the cumulative recruitment measure, and a joint sweep that probes practical identifiability.

The time-dependent recruitment rate function *α*(*t*) = *α*_0_ exp( −*t/τ*) has two parameters that jointly control the function’s shape: the initial recruitment amplitude *α*_0_ and the recruitment decay timescale *τ* . To assess how each parameter influences the forward model, we perform two controlled parametric sweeps, first varying *α*_0_ ∈ [0.4, 3.0] min^−1^ at fixed *τ* = 10 min (Figure 6, top), and then varying *τ* ∈ [2, 30] min at fixed *α*_0_ = 1.0 min^−1^ (Figure 6, bottom).

**Figure 6.**
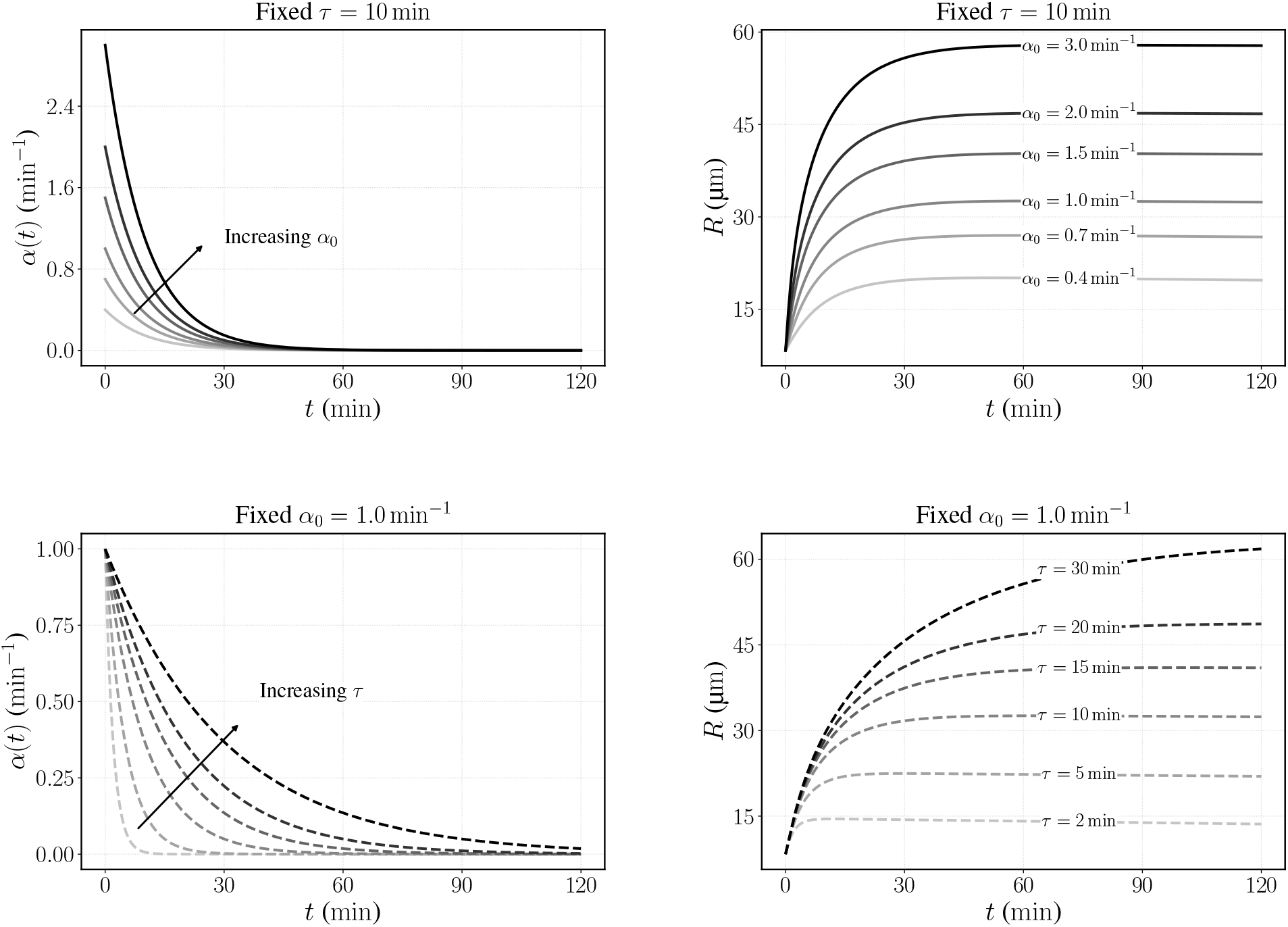
Parametric study showing the effect of varying initial recruitment amplitude *α*_0_ (top) and recruitment decay timescale *τ* (bottom) on the neutrophil recruitment dynamics *α*(*t*) (left) and swarm-front trajectories *R*(*t*) (right).

Increasing *α*_0_ raises the initial recruitment amplitude, producing a steeper early-time growth and a larger final swarm radius at fixed *τ* (Figure 6, top). Meanwhile, increasing *τ* prolongs the active recruitment window (Figure 6, bottom): by *t* = *τ* the recruitment rate has decayed to exp(−1) ≈ 37% of the initial value, and by *t* = 3*τ* recruitment is effectively shut off (exp(−3) ≈ 5%). A longer *τ* therefore allows the swarm to continue expanding for a longer duration, producing a larger terminal swarm radius. The observation that both *α*_0_ and *τ* affect the final swarm radius suggests that the parameters might not be uniquely determined. This motivates a closer look at how the recruitment rate function affects the swarming size.

The cumulative recruitment measure over an observation window [0, *t*_max_] is obtained by integrating Equation 2:

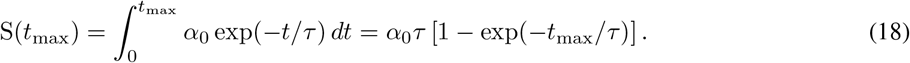

For a sufficiently long observation window relative to the timescale of recruitment decay, *t*_max_*/τ* ≫ 1, the exponential term becomes negligible and the integral approaches its asymptotic value

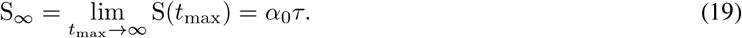

This asymptotic limit motivates the introduction of a derived dimensionless parameter

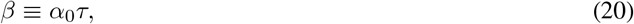

which represents the cumulative recruitment measure of the swarm. Together, the three quantities (*α*_0_, *τ, β*) characterize complementary aspects of the recruitment rate function: *α*_0_ controls the recruitment rate at *t* = 0, *τ* sets the timescale over which neutrophil sensitivity to LTB4 decays, and *β* approximates the cumulative recruitment of neutrophils over the duration of the swarm. However, only two of these parameters are independent. Because *α*_0_ and *β* capture clear, physically intuitive features of the swarm, here we choose to present our results in terms of them.

If *β* alone determines the cumulative recruitment of neutrophils, then trajectories with the same *β* but different *α*_0_ should converge to the same terminal swarm radius despite differing in their transient dynamics. To test this, we compute the final swarm radius for a range of parameters *α*_0_ ∈ [0.3, 2.0]min^−1^ and *β* ∈ [5, 50] (Figure 7). As expected, the final swarm radius is strongly related to *β* and nearly insensitive to *α*_0_. In the upper-left region, where *α*_0_ is low and *β* is high, the iso-boundary lines curve slightly; this is because in this regime the recruitment rate function has not fully decayed by *t*_max_, so the actual extent of recruitment differs from the asymptotic value.

**Figure 7.**
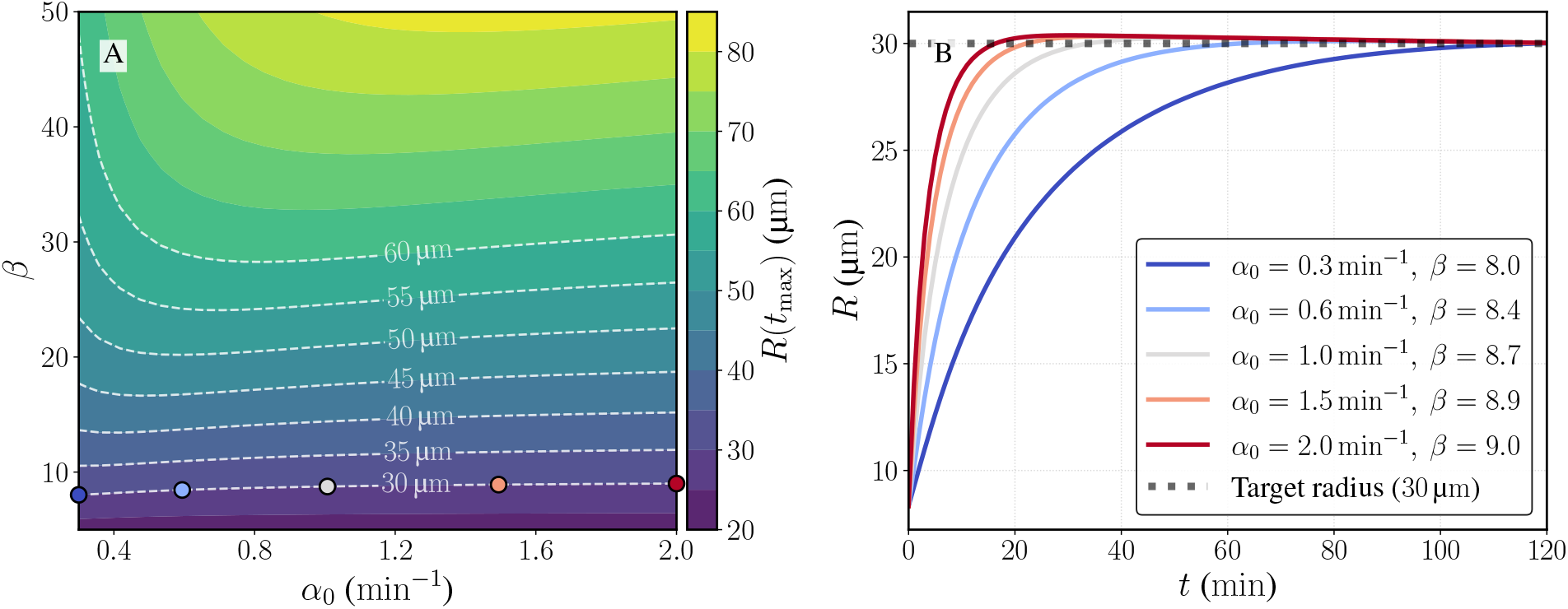
Identifiability analysis of the recruitment parameters. (A) Terminal swarm radius over a 30 *×* 30 grid in (*α*_0_, *β*). Dashed white iso-radius contours (30 to 60 µm, 5 µm spacing) run nearly parallel to the *α*_0_ axis, indicating that *β* predominantly controls the terminal swarm radius. (B) Swarm-front trajectories for five (*α*_0_, *β*) pairs sampled along the 30 µm iso-contour. All five reach the same terminal radius but differ in transient dynamics, demonstrating that the full trajectory determines parameters uniquely.

To illustrate identifiability explicitly, we sampled five distinct (*α*_0_, *β*) pairs along the 30 µm contour line and plotted their swarm-front trajectories (Figure 7B). All five trajectories converge to the same terminal radius of 30 µm at *t*_max_ = 120 min, confirming that the terminal swarm radius alone cannot uniquely determine the parameter pair. However, because the temporal dynamics vary significantly, uniqueness is recovered when information from multiple time points is used. Thus, our inverse problem satisfies the uniqueness criterion for a well-posed inverse problem.

Finally, we note that along a fixed-*β* contour, the trajectories become increasingly similar as *α*_0_ increases and *τ* = *β/α*_0_ decreases. Two challenges arise in this regime (Figure 7b). First, the practical identifiability of the inverse problem is reduced, and measurement noise, particularly in the earlier time points of larger *α*_0_ cases, will affect the inference. Second, the fast wavefront propagation during the initial recruitment time introduces numerical stiffness. For this reason, future experimental characterization of strongly swarming cases (larger *β*) should prioritize higher temporal resolution and measurement fidelity during the early swarming phase, where the trajectories carry the most discriminative information about (*α*_0_, *τ*). Furthermore, because the recruitment rate function is assumed to be a monotonically decreasing recruitment signal (Equation 2), the resulting inverse problem is therefore best suited for swarm trajectories that exhibit a smooth transition from early expansion to gradual deceleration and plateauing. More complex dynamics, such as delayed activation, secondary recruitment waves, or non-monotonic re-acceleration, would require a more flexible parameterization of *α*(*t*).

### 4.2 PINN prediction based on synthetic data

We validate our method by performing parameter inference on synthetic data (Figure 8). Both the data loss and the PDE residual loss decrease substantially during Adam optimization and are further refined by the subsequent L-BFGS step (Figure 8A). The predicted swarm-front trajectory closely matches the synthetic reference solution at the observation times *t*_*i*_ ∈ { 0, 10, 20, 30, 60, 90, 120} min (Figure 8B), demonstrating that the PINN captures the self-limiting dynamics of the swarm. Furthermore, both learned recruitment parameters, *α*_0_ and *τ*, converge to their ground-truth values after approximately 3 *×* 10^4^ training iterations (Figure 8C,D). Finally, the comparison between the predicted and reference radial density profiles (Figure 8E) further demonstrates that the trained PINN accurately reconstructs the full spatiotemporal cell density field. The relative *L*_2_ error, which is small across all validation time points, is calculated at each observation time as

**Figure 8.**
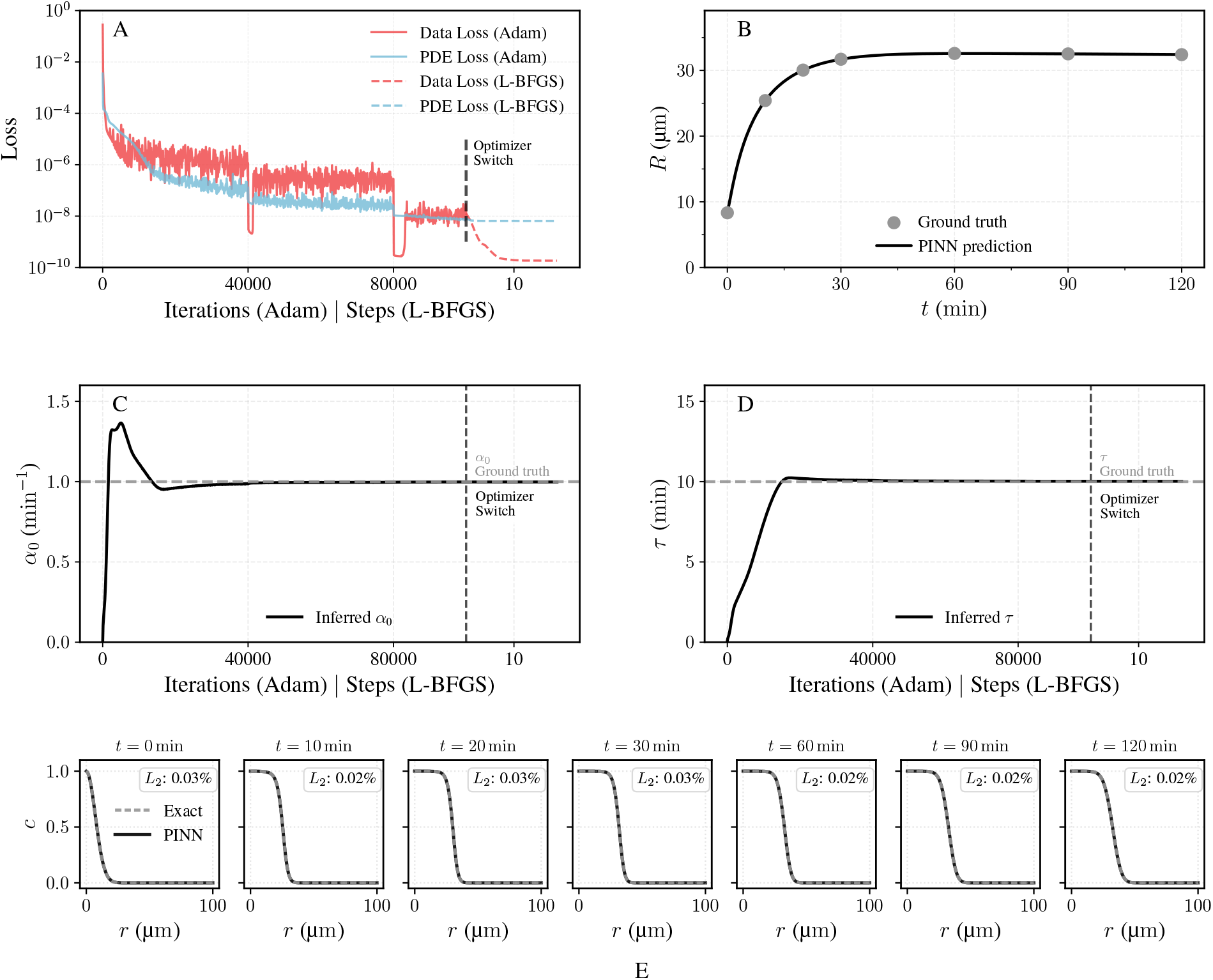
PINN results on synthetic data. (A) Evolution of the data loss and PDE residual loss during Adam optimization followed by L-BFGS refinement. (B) Predicted swarm-front trajectory compared with the synthetic reference solution. (C–D) Training trajectories of the inferred initial recruitment amplitude *α*_0_ and recruitment decay timescale *τ*, with dashed horizontal lines denoting the ground-truth values. (E) Predicted cell-density profiles compared with synthetic reference data at discrete time points. Differences are reported as relative *L*_2_ errors defined in Equation 21.

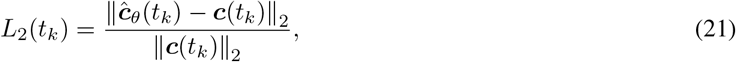

where in this validation case ***c***(*t*_*k*_) denotes the synthetic reference cell profile.

### 4.3 B-PINN prediction based on noisy synthetic data

Next, we perform uncertainty quantification with the B-PINN (Figure 9). The posterior mean closely matches the trajectory of both the noise-free synthetic data and the mean of the noisy synthetic data (Figure 9A). Compared with the broader prior predictive band, the posterior predictive band is substantially narrower and remains centered around the observed swarm trajectory, indicating that the noisy observations provide informative constraints on the governing dynamics. These results demonstrate that the B-PINN framework can recover the underlying self-limiting swarm evolution while quantifying uncertainty induced by noisy observations. In the joint posterior distribution *P*(*α*_0_, *τ* | ***c, R***), the HMC samples form a compact posterior cloud centered around the ground-truth crosshair 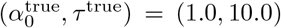 (Figure 9B). The posterior mean (*•*) lies close to the ground-truth crosshair, and the maximum a posteriori (MAP) estimate (⋆) falls within the high-density region of the posterior, providing a self-consistency check on the B-PINN forward model and the HMC sampler. We quantified parameter correlation by the Pearson correlation coefficient,

**Figure 9.**
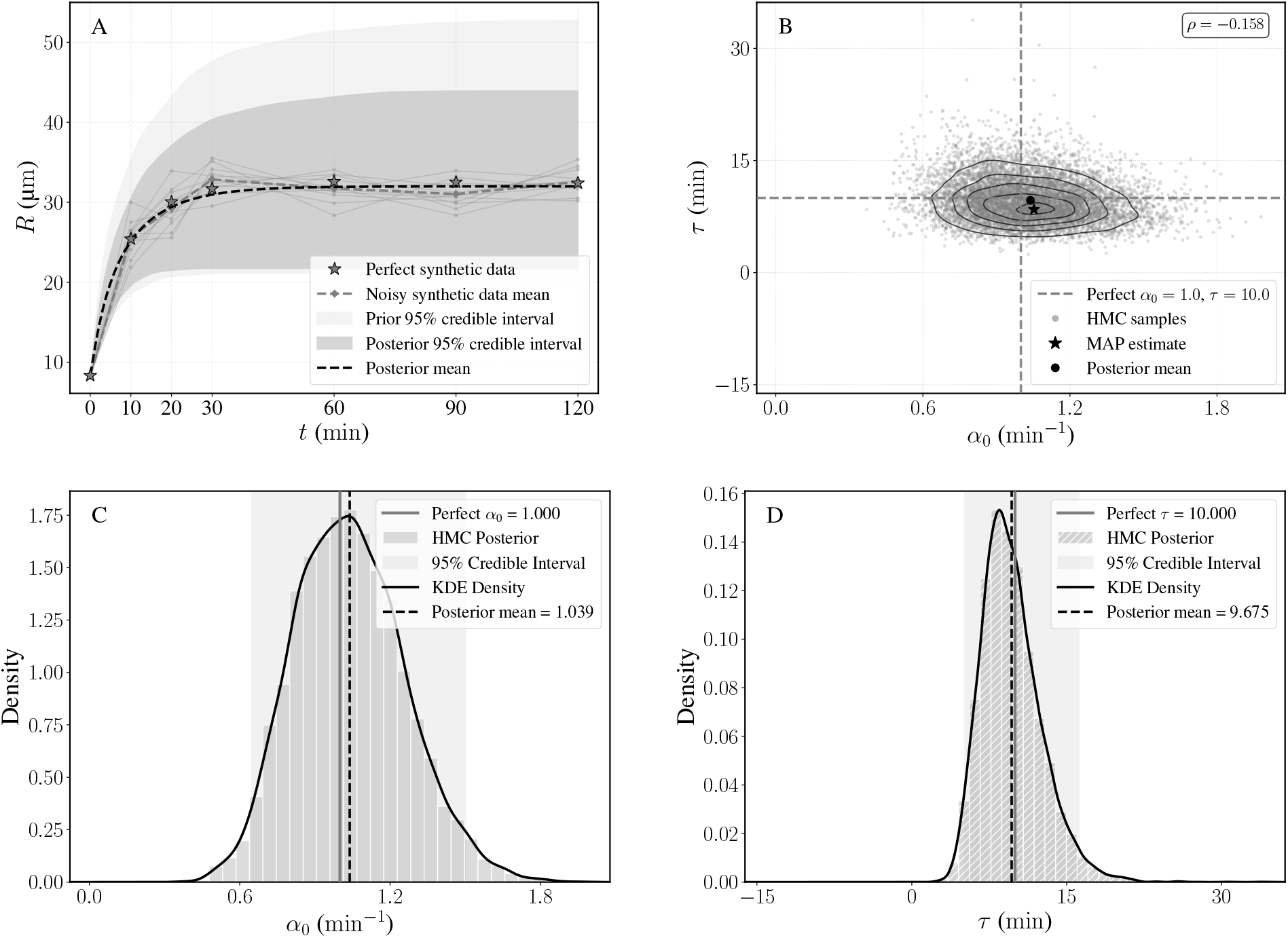
B-PINN prediction and uncertainty quantification on noisy synthetic data. (A) Swarm-front trajectory showing the noise-free synthetic reference solution, noisy synthetic observations, and the prior and posterior 95% bands. (B) Joint posterior distribution of *α*_0_ and *τ*, where points denote posterior samples, and contours show kernel density estimates (KDE) of the joint posterior density. (C–D) Marginal posterior distributions of *α*_0_ and *τ*, with 95% credible intervals and ground-truth values indicated.

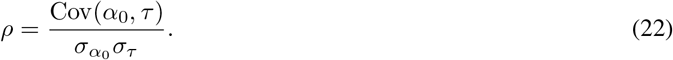

The weak (*ρ* = −0.158) correlation indicates only a mild trade-off in which a higher initial recruitment amplitude *α*_0_ can be partially compensated by a shorter recruitment decay timescale *τ* .

The marginal posterior distributions of *α*_0_ and *τ* (Figure 9C,D) are nearly centered about the ground truth. The *α*_0_ posterior is centered slightly above the ground truth and mildly right-skewed, whereas the *τ* posterior is centered slightly below the ground truth and has a more pronounced right tail. The broader relative spread of the *τ* posterior indicates that the recruitment decay timescale is less tightly constrained than the initial recruitment amplitude. The posterior means are *α*_0_ = 1.039 min^−1^ and *τ* = 9.675 min, compared with the ground-truth values *α*_0_ = 1.0 min^−1^ and *τ* = 10 min. The posterior means are within 5% in both cases, with relative errors of 3.90% for *α*_0_ and 3.25% for *τ* . These results demonstrate that the B-PINN framework recovers the governing recruitment parameters and their associated uncertainty from noisy observations.

### 4.4 PINN prediction based on the human and murine experimental data

The PINN was used to identify the recruitment parameters *α*_0_ and *τ* for each species/cluster-size condition. These inferred parameters were then substituted back into the governing equation, which was solved numerically using `SciPy` to obtain the swarm-front trajectory (Figure 10). Despite the sparsity and noise in the measurements, forward-evaluated trajectories closely reproduce the experimental swarm radii. In both species, the final swarm radius increases with HKSA bioparticle cluster size, indicating stronger neutrophil recruitment for larger stimuli.

**Figure 10.**
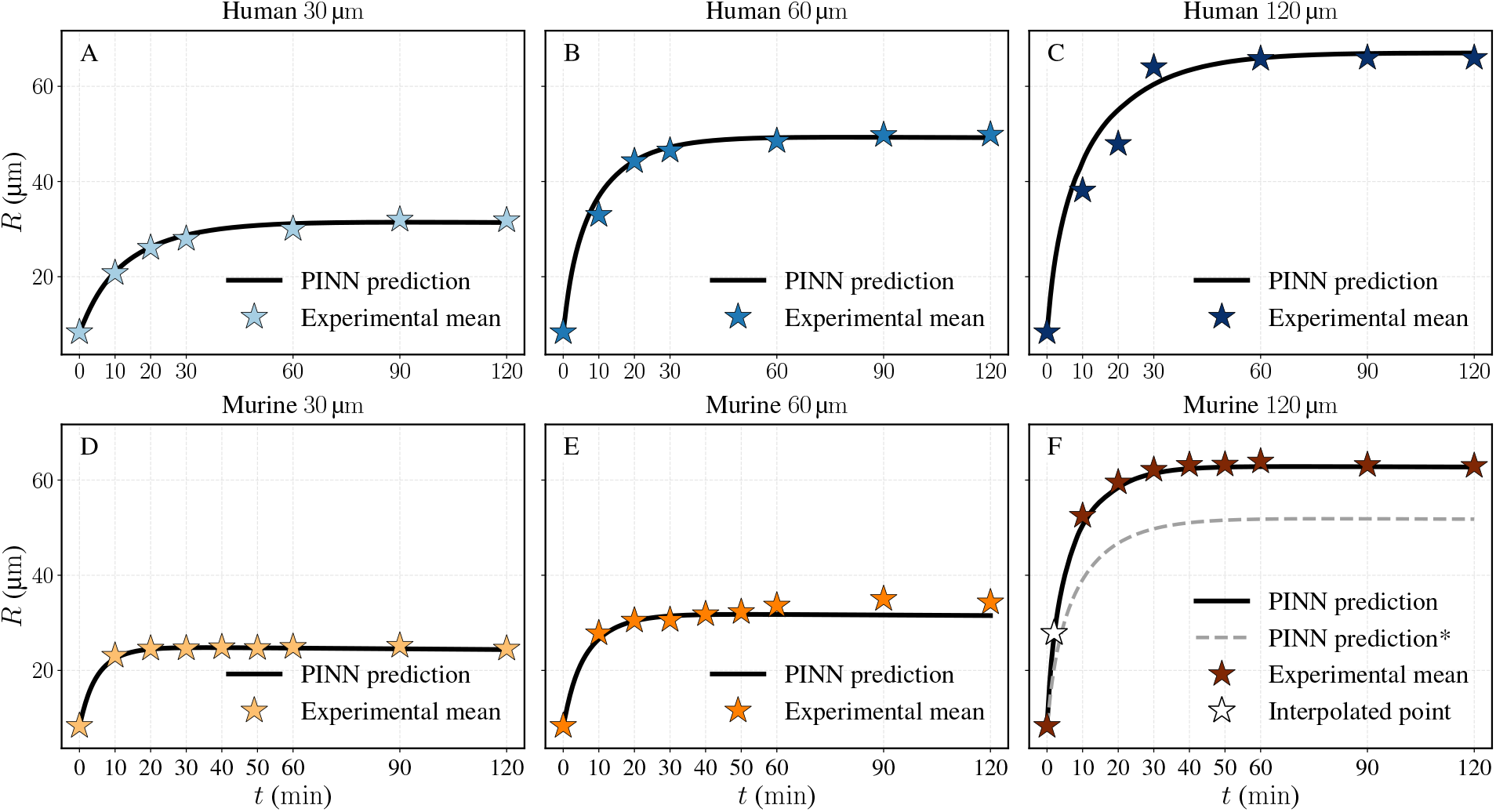
PINN-predicted swarm-front trajectories for human (A–C) and murine (D–F) neutrophils across the three HKSA bioparticle cluster-size conditions. Colored stars denote the mean experimental swarm-front radii, and solid black curves denote forward-solved trajectories from the PINN-inferred (*α*_0_, *τ* ). The open star in panel F marks the augmented *t* = 2 min early-time anchor point used for the murine 120 µm fit. The gray curve in panel F shows the corresponding PINN trajectory fit using only the raw experimental data, without the augmented early-time anchor. This unaugmented trajectory undershoots the experimental data points.

As discussed in subsection 4.1, our inverse modeling framework performs best when the swarm dynamics are smooth and monotonically decelerate toward a plateau. The 30 µm human and murine conditions follow this trend and can be fit without additional treatment, whereas the larger-cluster cases, particularly the 60 µm murine and 120 µm human datasets, exhibit transient non-monotonic wavefront dynamics (see Figure 4) that are only partially captured by the strictly decreasing recruitment rate function in Equation 2.

A separate numerical challenge occurs for the 120 µm murine case, which exhibits the strongest early swarming among all experimental cases. The rapid initial expansion produces a large time derivative *d****ĉ***_*θ*_*/dt*, which makes the optimization more sensitive to the PDE loss (Equation 12). In this regime, the sparse early-time observations provide limited constraint on the source term, and the PINN can underestimate the initial recruitment amplitude, leading to a systematic undershoot of the swarm-front trajectory (Figure 10F). To stabilize the fit during this early transient, we introduced a single augmented data point at *t* = 2 min to anchor the initial expansion. The augmented point was obtained by interpolation between the initial condition and the first measured time point and was used only for this condition.

For each experimental group (human and murine neutrophils responding to different bioparticle cluster sizes), the PINN infers values of *α*_0_ and *τ* . Because *α*_0_ controls the initial recruitment amplitude while *β* = *α*_0_*τ* concisely represents the cumulative recruitment measure, we calculate *β* from the inferred values and report them here for easier interpretation (Figure 11).

**Figure 11.**
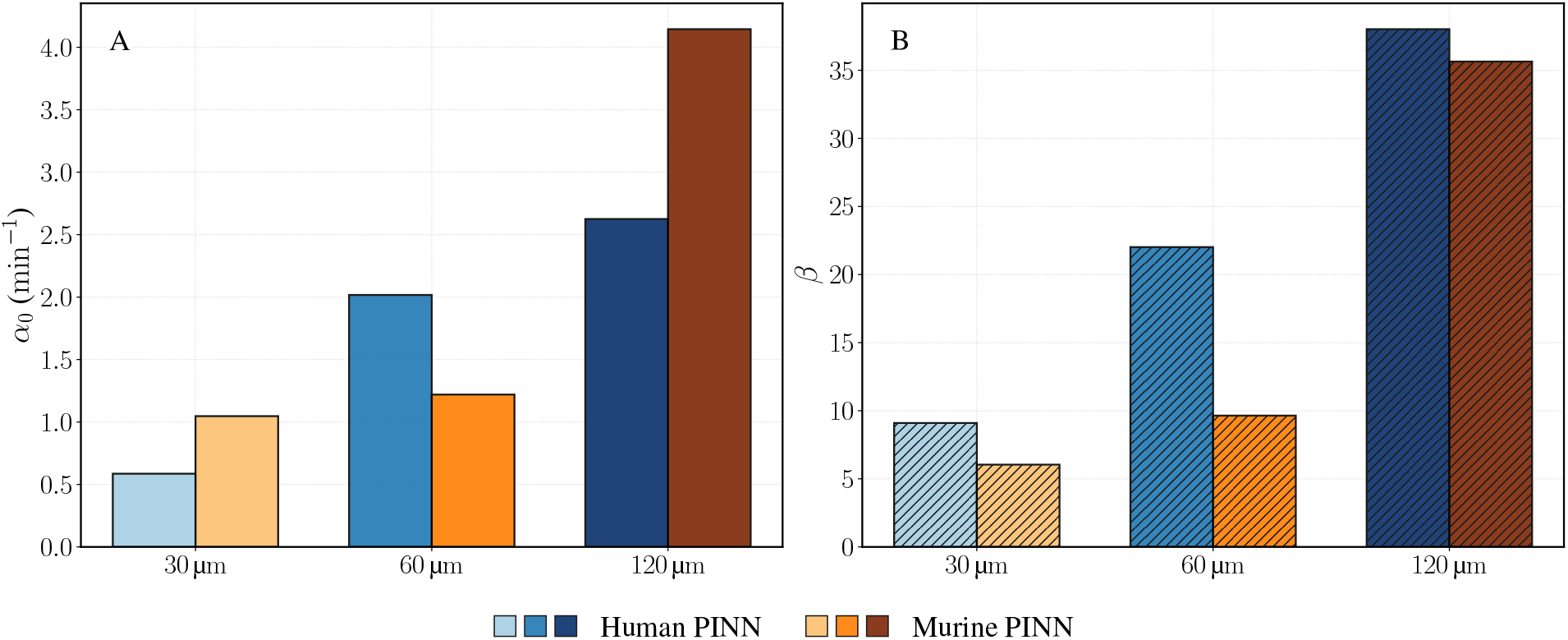
PINN estimates of initial recruitment amplitude *α*_0_ (A) and dimensionless cumulative recruitment measure *β* = *α*_0_*τ* (B), for human and murine neutrophil swarming across the three HKSA bioparticle cluster sizes.

Both species exhibit stronger initial recruitment (increasing *α*_0_) with increasing bioparticle cluster size, though the trend differs between species. In human neutrophils, *α*_0_ increases substantially from the 30 µm to the 60 µm case but shows diminishing gains for the 120 µm case. In contrast, murine neutrophils show that the change from 30 µm to 60 µm bioparticle cluster size is comparatively small, whereas 120 µm produces a pronounced rise, to the strongest initial recruitment amplitude among all cases.

Both species also show increasing cumulative recruitment measure (*β*) with increasing bioparticle cluster size. This trend is consistent with the larger final swarm radii observed for the larger bioparticle cluster size (Figure 10). Human neutrophils show an approximately 4.2-fold increase in *β* across the three bioparticle cluster sizes, whereas murine neutrophils show a nearly 6-fold increase. Notably, the murine 120 µm reaches a cumulative recruitment measure close to the human 120 µm, despite the different species-specific trends in *α*_0_.

### 4.5 B-PINN prediction based on the human and murine experimental data

The B-PINN generated posterior trajectories that closely reproduce the observed self-limiting swarm-front dynamics across the six experimental conditions (Figure 12). The posterior mean generally aligns with the experimental mean, and the 95% bands enclose the experimental data in nearly every case. While the inferred dynamics capture the overall swarm-front trajectory, the posterior mean shows a mild undershoot at later times.

**Figure 12.**
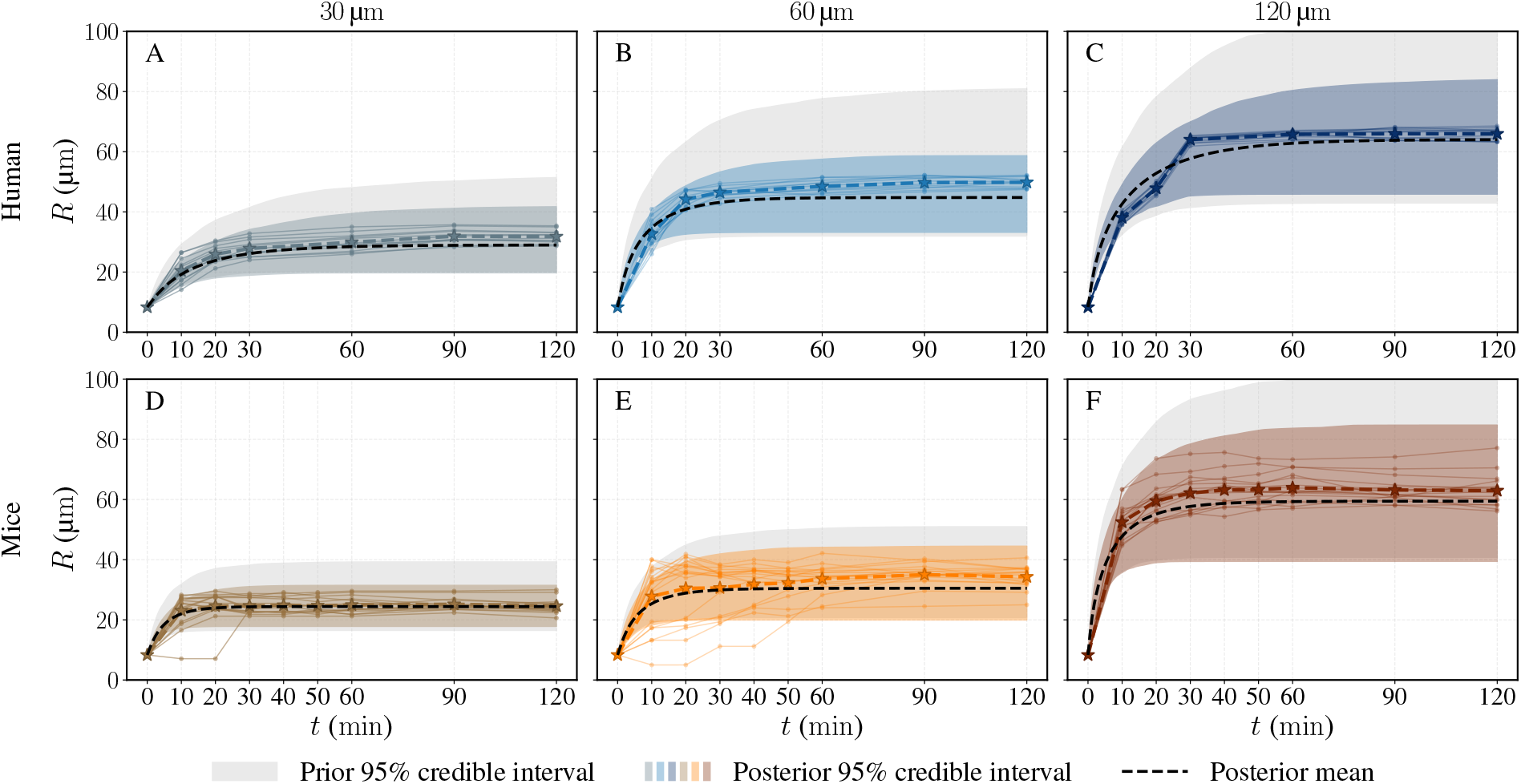
B-PINN posterior predictive swarm-front trajectories for human (A–C) and murine (D–F) neutrophils across the three HKSA bioparticle cluster sizes. Thin colored trajectories denote individual replicates, while the colored stars and dashed lines denote the replicate means. The gray shaded regions indicate the prior predictive 95% bands, the colored shaded regions indicate the posterior predictive 95% bands, and the black dashed lines show the posterior predictive means.

The posterior predictive bands are consistently narrower than the prior predictive bands, indicating that the experimental data provide substantial information for updating the PINN-based prior. Across conditions, the average reduction in 95% predictive interval width ranges from approximately 19% to 48%. This nonuniform contraction suggests that parameter identifiability depends on both the experimental condition and replicate variability.

The posterior distributions of the B-PINN-inferred recruitment parameters (Figure 13 and 14) quantify uncertainty in each parameter after conditioning on the experimental swarm-front radius data and the physics-informed priors. For all six experimental conditions, the posterior means lie close to the deterministic PINN estimates of both *α*_0_ and *τ* . Thus, the PINN estimates remain compatible with the Bayesian posterior.

**Figure 13.**
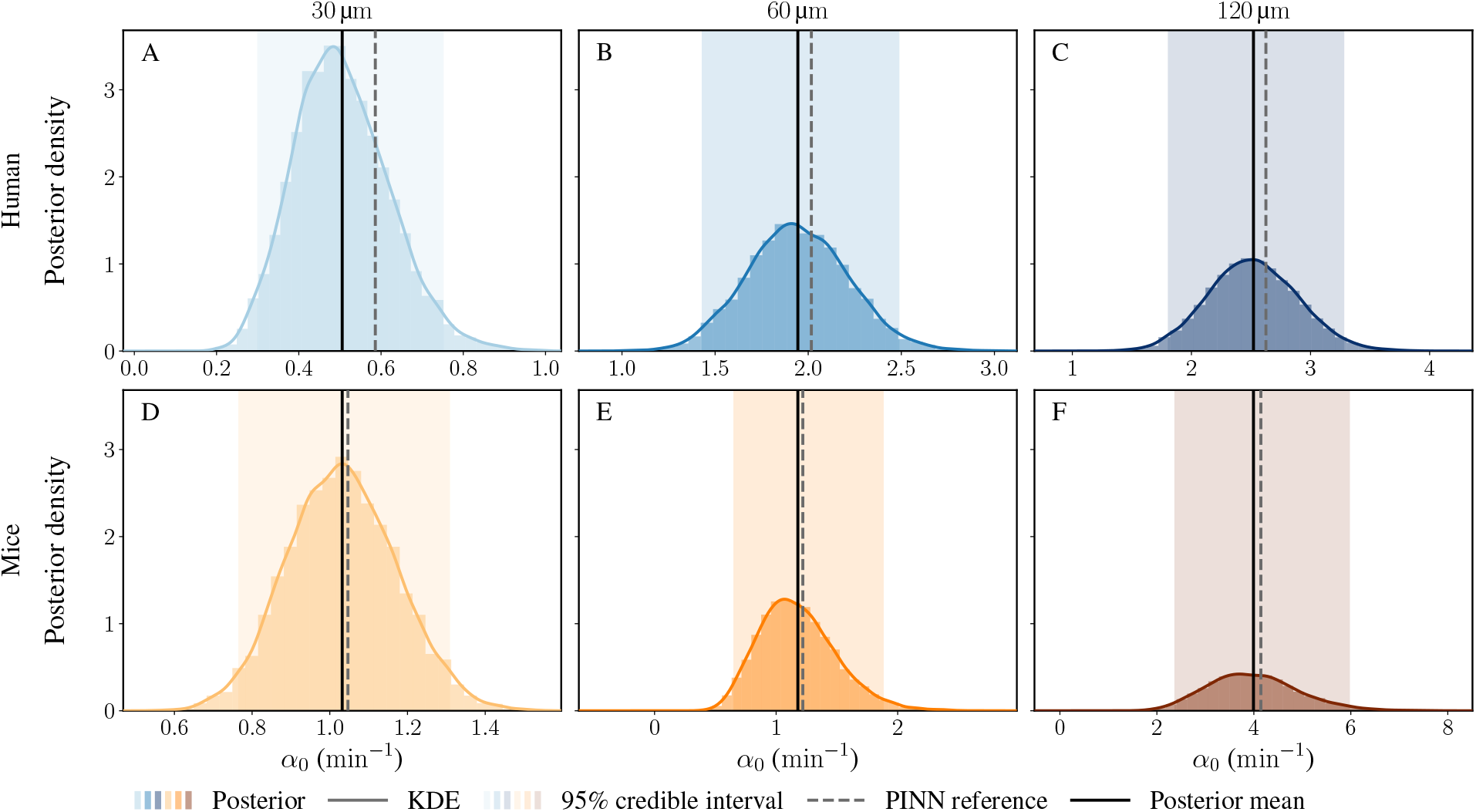
B-PINN inference of the initial recruitment amplitude *α*_0_ for human (A–C) and murine (D–F) neutrophils across the three HKSA bioparticle cluster-size conditions. Histograms denote HMC posterior samples, solid colored curves show kernel density estimates, and shaded regions indicate 95% credible intervals. Dashed vertical lines mark the PINN-inferred values of *α*_0_, while solid black vertical lines mark the B-PINN posterior means.

**Figure 14.**
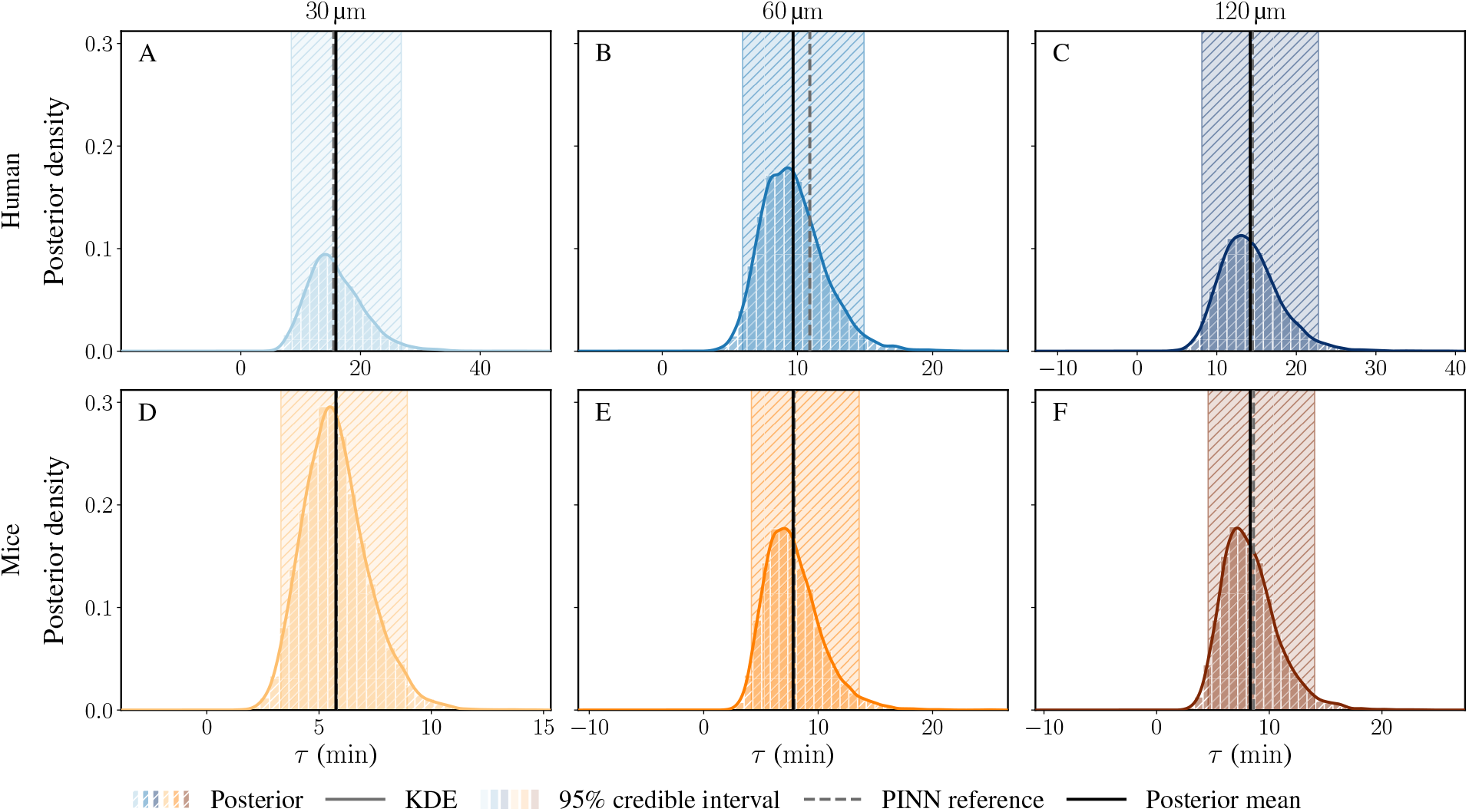
B-PINN inference of the recruitment decay timescale *τ* for human (A–C) and murine (D–F) neutrophils across the three HKSA bioparticle cluster-size conditions. Histograms denote HMC posterior samples, solid colored curves show kernel density estimates, and shaded regions indicate 95% credible intervals. Dashed vertical lines mark the PINN-inferred values of *τ*, while solid black vertical lines mark the B-PINN posterior means.

For *α*_0_ (Figure 13), the posterior means are consistently slightly lower than the PINN point estimates, with an average decrease of approximately 5% and a largest shift of 13.7% in the human 30 µm condition. This modest downward shift is consistent with conditioning on replicate variability and the physics-residual likelihood, which favor smoother posterior trajectories when temporal observations are sparse. The recruitment decay timescale *τ* (Figure 14) shifts even less (an average decrease of approximately 2.6%), consistent with the mild coupling between *α*_0_ and *τ* .

To provide a complementary view of parameter uncertainty and coupling, we examined the joint posterior distributions of *α*_0_ and *τ* (Figure 15). Across all six conditions, the posterior clouds and density contours are centered near the deterministic PINN estimates. This confirms that the PINN solutions are compatible with Bayesian inference while also revealing the range of parameter pairs supported by the data and physics-informed likelihood.

**Figure 15.**
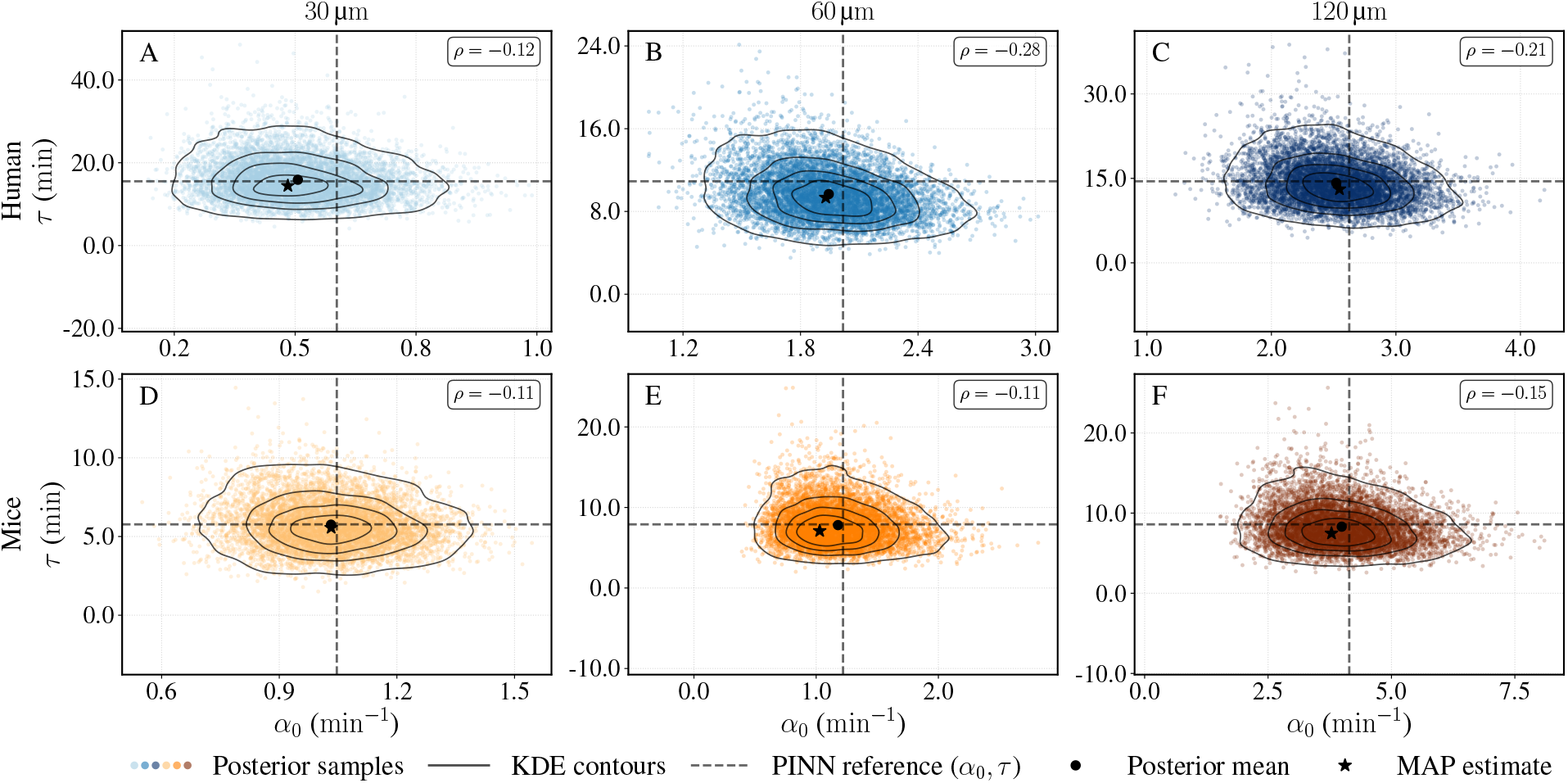
Joint posterior distributions of *α*_0_ and *τ* for human (A–C) and murine (D–F) neutrophils exposed to HKSA bioparticle clusters of different diameters. Points denote HMC posterior samples, and black contours show kernel-density estimates of the joint posterior density. The black dashed lines represent the deterministic PINN estimates of *α*_0_ and *τ* for each condition.

The joint posteriors exhibit weak negative correlations between *α*_0_ and *τ*, with Pearson correlation coefficients ranging from approximately −0.28 to −0.11 across conditions. This indicates a partial trade-off between the initial recruitment amplitude and the recruitment decay timescale: a larger *α*_0_ can be partially compensated by a shorter *τ* while preserving a similar swarm-front trajectory. However, the correlation is not strong enough to indicate a severe non-identifiability ridge. This partial coupling further motivates using the derived quantity *β* = *α*_0_*τ* as a complementary summary.

The B-PINN posterior quantifies uncertainty in the inferred recruitment parameters (Figure 16). Blue and orange brackets show adjacent-size posterior contrasts within the human and murine groups, respectively, whereas black brackets show human-murine contrasts at the same bioparticle cluster sizes. Only contrasts satisfying *p*_tail_ *<* 0.05 are shown, unresolved adjacent comparisons are left unmarked.

**Figure 16.**
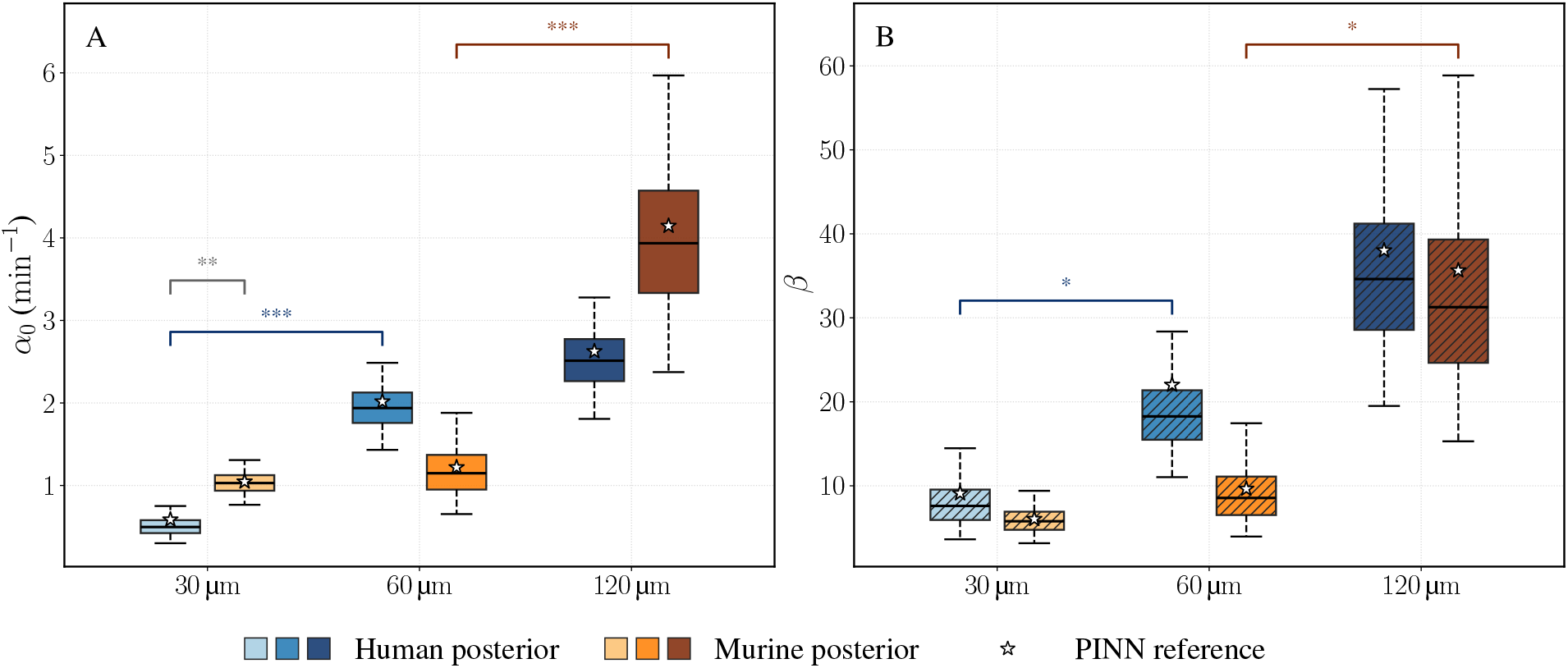
B-PINN posterior distributions of the initial recruitment amplitude *α*_0_ (A) and cumulative recruitment measure *β* = *α*_0_*τ* (B) for human and murine neutrophils across HKSA bioparticle cluster sizes. Boxes span the interquartile ranges of the posterior HMC samples, horizontal lines indicate posterior medians, whiskers mark the 2.5th and 97.5th percentiles (95% credible intervals). White star markers denote the deterministic PINN estimates. Colored brackets compare adjacent bioparticle cluster sizes within the same species, whereas black brackets compare human and murine posteriors at the same bioparticle cluster sizes. Asterisks indicate two-sided posterior tail probabilities: * for *p*_tail_ *<* 0.05, ** for *p*_tail_ *<* 0.01, and *** for *p*_tail_ *<* 0.001. Comparisons with *p*_tail_ ≥ 0.05 are not shown. These annotations summarize Bayesian posterior contrasts and are not frequentist *t*-test *p*-values.

For the initial recruitment amplitude *α*_0_, the human posterior medians (posterior standard deviations) were 0.50 (0.12), 1.94 (0.27), and 2.51 (0.38) min^−1^ at 30 µm, 60 µm, and 120 µm, respectively. The corresponding murine values were 1.03 (0.14), 1.15 (0.31), and 3.94 (0.92) min^−1^. The adjacent-size contrasts resolved the increase from 30 µm to 60 µm in human neutrophils and from 60 µm to 120 µm in murine neutrophils, but not the remaining two comparisons. Among the same size human and murine comparisons, a species difference was resolved only at 30 µm (*p*_tail_ = 0.0046). The corresponding species contrasts at 60 µm and 120 µm were not resolved. Additional independent replicates and early-time measurements would therefore be most informative for differentiating *α*_0_ between the human 60 µm and 120 µm conditions and between the murine 30 µm and 60 µm conditions.

The posterior medians of the cumulative recruitment measure *β* = *α*_0_*τ* increased with bioparticle cluster size in both species. The human posterior medians (posterior standard deviations) were 7.61 (2.79), 18.27 (4.44), and 34.62 (9.74), while the corresponding murine values were 5.78 (1.60), 8.56 (3.53), and 31.27 (11.35). From 30 µm to 120 µm, the median increased by 355% in human neutrophils and by 441% in murine neutrophils. Among the shown adjacent-size comparisons, the human 30 µm-to-60 µm increase and the murine 60 µm-to-120 µm increase were resolved, whereas the other two comparisons were not. The 30 µm-to-120 µm posterior contrasts also supported an overall increase in both species, although these nonadjacent comparisons are not shown.

The matched-size human-murine contrasts did not meet the *p*_tail_ *<* 0.05 criterion for *β* at any bioparticle cluster size. At 120 µm, the human and murine posterior medians were similar (34.62 and 31.27, respectively) relative to their uncertainty. This result is consistent with comparable cumulative recruitment at the largest bioparticle cluster size, but it does not imply identical temporal recruitment dynamics because the same *β* can arise from different combinations of *α*_0_ and *τ* (subsection 4.1).

## 5 Future directions

The present study establishes an interpretable framework for inferring self-limiting neutrophil recruitment from sparse experimental observations and provides several directions for further development. One important direction is to extend the phenomenological Fisher-KPP model toward a more mechanistic description of neutrophil signaling and migration. In the current formulation, recruitment attenuation is represented by the exponentially decaying recruitment rate function *α*(*t*). A coupled PDE system describing chemoattractant transport, neutrophil migration, and receptor desensitization could instead allow swarm initiation and shutdown to emerge from the modeled interactions. More flexible or data-driven forms of *α*(*t*) could also represent delayed activation, transient recruitment peaks, and secondary recruitment waves. A second direction is to infer the initial neutrophil density profile directly from early-time experimental images. The smooth Gaussian profile used here provides a numerically stable and consistent initial condition across experimental conditions, but its prescribed shape can influence the inferred values of *α*_0_ and *τ* . Experimentally, denser temporal measurements during the early expansion phase would also improve the identifiability of these two kinetic parameters. A third direction is to extend the radially symmetric formulation to two-dimensional, image-informed cell density fields. Radial symmetry provides an effective description of the mean expansion of approximately circular swarms, whereas a two-dimensional model could capture asymmetric spreading, branching, local variations in cell density, and heterogeneous signaling or tissue environments. Such an extension enable the framework to model neutrophil swarming in more complex *in vivo* settings. Together, these extensions would advance the framework from an effective description of swarm expansion toward a spatially resolved, mechanistic, and predictive model of neutrophil collective behavior.

## 6 Conclusion

Neutrophils are an essential component of innate immunity, providing rapid, non-specific defense despite substantial heterogeneity in maturation state, tissue location, activation status, and disease context. Although neutrophil swarming exhibits conserved qualitative features across species, its quantitative dynamics differ across commonly studied experimental systems, including zebrafish, murine, and human neutrophils. In this work, we developed deterministic and Bayesian physics-informed neural network models to infer effective recruitment dynamics from human and murine neutrophil swarming data. The PINN identified species- and cluster-size-dependent recruitment parameters, while the B-PINN extended these estimates by quantifying posterior uncertainty and revealing which parameter trends were robust under sparse, noisy swarm-front radius observations. The results show that larger HKSA bioparticle clusters generally yield higher values of the cumulative recruitment measure. Specifically, as the HKSA bioparticle cluster diameter increased from 30 µm to 120 µm, the cumulative recruitment measure increased approximately 4.2-fold for human neutrophils and nearly 6-fold for murine neutrophils. At the same time, the B-PINN posterior distributions highlight important limitations: the inferred *α*_0_ and *τ* values are weakly coupled through accumulated recruitment dynamics, and the posterior predictive trajectories show mild late-time undershoot relative to experimental swarm radius data. These findings indicate that additional measurements during the early, fast-expanding phase of swarming would improve parameter identifiability and biological interpretability. Overall, this modeling framework provides a quantitative bridge for comparing murine and human neutrophil swarming and offers a path toward more mechanistic translations of preclinical immune-dynamics data.

## 7 Acknowledgments

Computational resources were provided in part by the Notre Dame Center for Research Computing. The authors also thank Dr. Dodi Heryadi for assistance with software and computational resources, and Dr. Evelyn Strickland, Dr. Pan Deng, and Dr. Zhen Zhang for helpful discussions.

## Data Availability Statement

All source code, synthetic datasets, trained model parameters, and processed experimental data supporting this study are openly available in the GitHub repository https://github.com/theCoMMaNDlab/neutrophil_swarming. The repository includes the PINN and B-PINN implementations, the numerical PDE solver and synthetic data-generation scripts, the processed swarm-front datasets, and the notebooks used to generate all figures in this manuscript. The underlying human and murine neutrophil cell-count measurements were originally reported by Glaser et al. [19]; the processed swarm-front values derived from them are provided in the repository.

